# Mega-scale movie-fields in the mouse visuo-hippocampal network

**DOI:** 10.1101/2022.12.07.519455

**Authors:** Chinmay S. Purandare, Mayank R. Mehta

## Abstract

Natural experience often involves a continuous series of related images while the subject is immobile. How does the cortico-hippocampal circuit process this information? The hippocampus is crucial for episodic memory^1–3^, but most rodent single unit studies require spatial exploration^4–6^ or active engagement^7^. Hence, we investigated neural responses to a silent, isoluminant, black and white movie in head-fixed mice without any task or locomotion demands, or rewards, from the Allen Brain Observatory. The activity of most neurons (97%, 6554/6785) in the thalamo-cortical visual areas was significantly modulated by the 30s long movie clip. Surprisingly, a third (33%, 3379/10263) of hippocampal –dentate gyrus, CA1 and subiculum– neurons showed movie-selectivity, with elevated firing in specific movie sub-segments, termed movie-fields. Movie-tuning remained intact when mice were immobile or ran spontaneously. On average, a tuned cell had more than 5 movie-fields in visual areas, but only 2 in hippocampal areas. The movie-field durations in all brain regions spanned an unprecedented 1000-fold range: from 0.02s to 20s, termed mega-scale coding. Yet, the total duration of all the movie-fields of a cell was comparable across neurons and brain regions. We hypothesize that hippocampal responses show greater continuous-sequence encoding than visual areas, as evidenced by fewer and broader movie-fields than in visual areas. Consistent with this hypothesis, repeated presentation of the movie images in a fixed, scrambled sequence virtually abolished hippocampal but not visual-cortical selectivity. The enhancement of continuous movie tuning compared to the scrambled sequence was eight-fold greater in hippocampal than visual areas, further supporting episodic-sequence encoding. Thus, all mouse-brain areas investigated encoded segments of the movie. Similar results are likely to hold in primates and humans. Hence, movies could provide a unified way to probe neural mechanisms of episodic information processing and memory, even in immobile subjects, across brain regions, and species.

## Introduction

In addition to the position and orientation of simple visual cues, like Gabor patches and drifting gratings^8^, primary visual cortical responses are also direction selective^9^, and show predictive coding^10^, suggesting that the temporal sequence of visual cues influences neural firing. Accordingly, these and higher visual cortical neurons too encode a sequence of visual images, i.e., a movie^11–18^. The hippocampus is farthest downstream from the retina in the visual circuit. The rodent hippocampal place cells encode spatial or temporal sequences^2,19–26^ and episode-like responses^27–29^. However, these responses typically require active locomotion^30^, and they are thought to be non-sensory responses^31^. Primate and human hippocampal responses are selective to specific sets of visual cues, e.g., the object-place association^32^, their short-term^1^ and longterm^33^ memories, cognitive boundaries between episodic movies^34^, and event integration for narrative association^35^. However, despite strong evidence for the role of hippocampus in episodic memory, the hippocampal encoding of a continuous sequence of images, i.e., a visual episode, is unknown.

## Results

### Significant movie tuning across cortico-hippocampal areas

We used a publicly available dataset (Allen Brain Observatory – Neuropixels Visual Coding, © 2019 Allen Institute). Mice were monocularly shown a 30s clip of a continuous segment from the movie *Touch of Evil* (Welles, 1958)^36^ (Figure 1-figure supplement 1 and Figure 1-Video 1). Mice were head-fixed but were free to run on a circular disk. A total of 17048 broad spiking, active, putatively excitatory neurons were analyzed, recorded using 4-6 Neuropixel probes in 24 sessions from 24 mice (See *Methods*).

**Figure 1.**
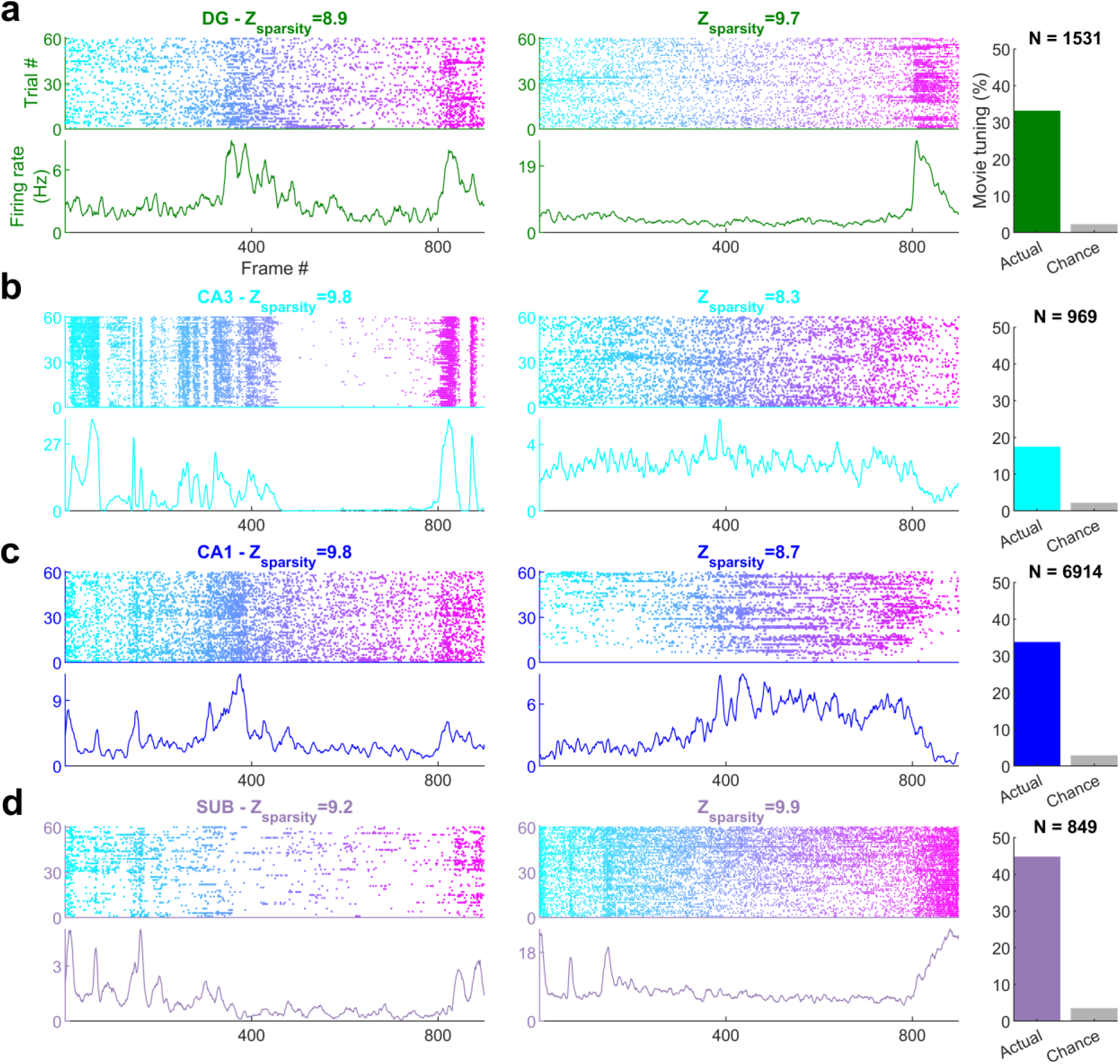
| Movie frame selectivity in hippocampal neurons (a) Raster plots of two different dentate gyrus (DG) neurons as a function of the movie frame (top) over 60 trials, and the corresponding mean firing rate response (bottom). These two cells had significantly increased activity in specific segments of the movie. Z-scored sparsity indicating strength of modulation is indicated above. 33.1% of dentate neurons were significantly modulated by the movie (right, green bar), far greater than chance (gray bar). Total active, broad spiking neurons for each brain region indicated at top (N_tuned_ /N_cells_=506/1531). **(b)** Same as (a), for CA3 (168/969, 17.3%), **(c)** CA1 (2326/6914, 33.6%) and **(d)** subiculum (379/849, 44.6%) neurons.

The majority of neurons in the visual areas (Lateral geniculate nucleus LGN, primary visual cortex V1, higher visual areas: antero-medial and posterior-medial AM-PM) were modulated by the movie, consistent with previous reports (Figure 1-figure supplement 2)^11–18^. Surprisingly, neurons from all parts of the hippocampus (dentate gyrus DG, CA3, CA1, subiculum SUB) were also clearly modulated (Figure 1), with reliable, elevated spiking across many trials in small movie segments. To quantify selectivity in an identical, firing rate- and threshold-independent fashion across brain regions, we computed the z-scored sparsity^37–40^ of neural selectivity (See *Methods*). Cells with z-scored sparsity >2 were considered significantly (*p*<0.03) modulated. Other metrics of selectivity, like depth of modulation or mutual information, provided qualitatively similar results (Figure 1-figure supplement 3). The areas V1 (97.3%) and AM-PM (97.1%) had the largest percentage of movie tuned cells. Similarly, the majority of neurons in LGN (89.2%) too showed significant modulation by the movie. This level of selectivity is much higher than reported earlier^11^ (∼40%), perhaps because we analyzed extracellular spikes, while the previous study used calcium imaging. On the other hand, the movie selectivity was greater than the selectivity for classic stimuli, like drifting gratings, in V1, even within calcium imaging data, in agreement of reports of better model fit with natural stimuli for primate visual responses^41^. Direct quantitative comparison across stimuli is difficult and beyond the scope of this study because the movie frames appeared every 30ms, and were preceded by similar images, while classic stimuli were presented for 250ms, in a random order. Thus, the vast majority of thalamo-cortical neurons were significantly modulated by the movie.

Movie selectivity was prevalent in the hippocampal regions too, despite head fixation, dissociation between self-movements and visual cues as well as the absence of rewards, task, or memory demands (Figure 1a-d). Subiculum, the output region of the hippocampus, farthest removed from the retina, had the largest fraction (44.6% Figure 1d) of movie-tuned neurons, followed by the upstream CA1 (33.6%, Figure 1c) and dentate gyrus (33.1%, Figure 1a). However, CA3 movie selectivity was nearly half as much (17.3%, Figure 1b). This is unlike place cells, where CA3 and CA1 selectivity are comparable^42,43^ and subiculum selectivity is weaker^44^.

### Movie tuning is not an artifact of behavioral or brain state changes

To confirm these findings, we performed several controls. Running alters neural activity in visual areas^45–48^ and hippocampus^49–51^. Hence, we used the data from only the stationary epochs (see *Methods*) and only from sessions with at least 300 seconds of stationary data (17 sessions, 24906 cells). Movie tuning was unchanged in this data (Figure 1-figure supplement 4). This is unlike place cells where spatial selectivity is greatly reduced during immobility^5,6^. Neurons recorded simultaneously from the same brain region also showed different selectivity patterns (Figure 1-figure supplement 5). Thus, nonspecific effects such as running cannot explain brain wide movie selectivity. Prolonged immobility could change the brain state, e.g., the emergence of sharp-wave ripples. Hence, we removed the data around sharp wave ripples and confirmed that movie tuning was unaffected (Figure 1-figure supplement 6). Strong movie tuned cells were seen in sessions with long bouts of running as well as with predominantly immobile behavior (Figure 1-figure supplement 7), unlike responses to auditory tones, which were lost during running behavior^51^. Place cell selectivity of hippocampal neurons is influenced by theta rhythm^52–54^. We compared the movie selectivity during periods of high theta, vs. periods of low theta. Significant movie selectivity in both cases (Figure 1-figure supplement 7). To further assess the effect of changes in brain state, we similarly analyzed movie tuning in two equal subsegments of data, corresponding to epochs with high and low pupil dilation, which is a strong correlate of arousal^55–57^. Movie tuning was above chance levels in both sub-segments (Figure 1-figure supplement 7). Hence, locomotion, arousal or changes in brain states cannot explain the hippocampal movie tuning.

### Similarities and differences between place fields and movie fields

Hippocampal neurons have one or two place fields in typical mazes which take a few seconds to traverse^58^. In larger arenas that take tens of seconds to traverse, the number of peaks per cell and the peak duration increases^59–62^. Peak detection for movie tuning is nontrivial because neurons have nonzero background firing rates, and the elevated rates cover a wide range (Figure 1). We developed a novel algorithm to address this (see *Methods*). On average, V1 neurons had the largest number of movie-fields (Figure 2a, mean±s.e.m.=10.4±0.1, here we use mean instead of median to gain a better resolution for the small and discrete values of number of fields per cell), followed by LGN (8.6±0.3) and AM-PM (6.3±0.07). Hippocampal areas had significantly fewer movie-fields per cell: dentate gyrus (2.1±0.1), CA3 (2.8±0.3), CA1(2.0±0.02) and subiculum (2.1±0.05). Thus, the number of movie-fields per cell was smaller than the number of place-fields per cell in comparably long spatial tracks^59–64^, but a handful of hippocampal cells had more than 5 movie-fields (Figure 2-figure supplement 1).

**Figure 2.**
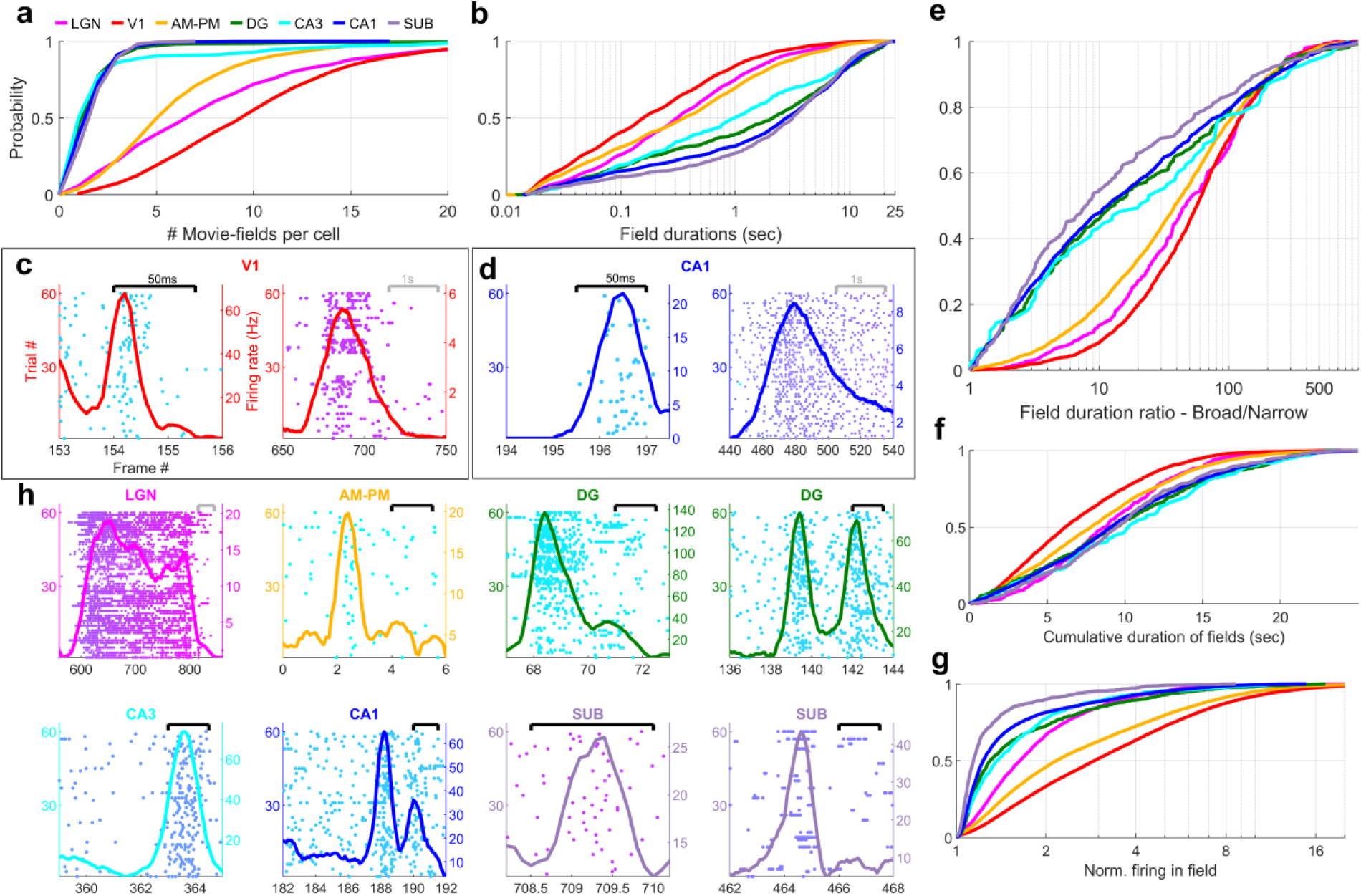
| Multi-peaked, mega-scale movie-fields across all brain areas (a) Distribution of the number of movie-fields per tuned cell (See *Methods)* in different brain regions (shown by different colors, top line inset, arranged in their hierarchical order). Hippocampal regions (blue-green shades) were significantly different from each other (KS-test *p*<0.04), except DG-CA3. All visual regions were significantly different from each other (KS-test p<7.0×10^-11^). All visual-hippocampal region pairwise comparisons were also significantly different (KS-test *p*<1.8×10^-44^). CA1 had the lowest number of movie-fields per cell (2.0±0.02, mean±s.e.m.) while V1 had the highest (10.4±0.1). **(b)** Distribution of the durations of movie-fields identified in (a), across all tuned neurons from a given brain region. These were significantly different for all brain region pairs (KS-test *p*<7.3×10^-3^). The longest movie-fields were in subiculum (3169.9±169.8 ms), and the shortest in V1 (156.6±9.2ms). **(c)** Snippets of movie-fields from an example cell from V1, with 2 of the fields zoomed in, showing 60x difference in duration. Black bar at top indicates 50ms, and gray bar indicates 1s. Each frame corresponds to 33.3ms. Average response (solid trace, y-axis on the right) is superimposed on the trial wise spiking response (dots, y-axis on the left). Color of dots corresponds to frame numbers as in Figure 1. **(d)** Same as (c), for a CA1 neuron with 54x difference in duration. **(e)** The ratio of longest to shortest field duration within a single cell, i.e., mega-scale index, was largest in V1 (56.7±2.2) and least in subiculum (8.0±9.7). All visual-visual and visual-hippocampal brain region pairs were significantly different on this metric (KS-test *p*<0.02). Among the hippocampal-hippocampal pairs, only CA3-SUB were significantly different (*p*=0.03). **(f)** For each cell, the total duration of all movie-fields, i.e., cumulative duration of significantly elevated activity, was comparable across brain regions. The largest cumulative duration (10.2±0.46s, CA3) was only 1.66x of the smallest (6.2±0.09 sec, V1). Visual-hippocampal and visual-visual brain region pairs’ cumulative duration distributions were significantly different (KS-test *p*<0.001), but not hippocampal pairs (*p*>0.07). **(g)** Distribution of the firing within fields, normalized by that in the shuffle response. All fields from all tuned neurons in a brain region were used. Firing in movie-fields was significantly different across all brain region pairs (KS-test, *p*<1.0×10^-7^), except DG-CA3. Movie-field firing was largest in V1 (2.9±0.03) and smallest in subiculum (1.14±0.03). **(h)** Snippets of movie-fields from representative tuned cells, from LGN showing a long movie-field (233 frames, or 7.8s, panel 1), and from AM-PM and from hippocampus showing short fields (2 frames or 66.6ms wide or less).

### Mega-scale structure of movie-fields

Typical receptive field size increases as one moves away from the retina in the visual hierarchy^36^. A similar effect was seen for movie-field durations. On average, hippocampal movie-fields were longer than visual regions (Figure 2b). But there were many exceptions –movie-fields of LGN (median±s.e.m., here and subsequently, unless stated otherwise, 308.5±33.9 ms) were twice as long as in V1 (156.6±9.2ms). Movie-fields of subiculum (3169.9±169.8 ms) were significantly longer than CA1 (2786.1±77.5 ms) and nearly three-fold longer than the upstream CA3 (979.1±241.1 ms). However, the dentate movie-fields (2113.2±172.4 ms) were two-fold longer than the downstream CA3. This is similar to the patterns reported for CA3, CA1 and dentate gyrus place cells^64^. But others have claimed that CA3 place fields are slightly bigger than CA1^65^, whereas movie-fields showed the opposite pattern.

The movie-field durations spanned a 500-1000-fold range in every brain region investigated (Figure 2e). This mega-scale scale is unprecedentedly large, nearly 2 orders of magnitude greater than previous reports in place cells^59,61^. Even individual neurons showed 100-fold mega-scale responses (Figure 2c & d) compared to less than 10-fold scale within single place cells^59,61^. The mega-scale tuning within a neuron was largest in V1 and smallest in subiculum (Figure 2e). This is partly because the short duration movie-fields in hippocampal regions were typically neither as narrow nor as prominent as in the visual areas (Figure 2-figure supplement 2).

Despite these differences in mega-scale tuning across different brain areas, the total duration of elevated activity, i.e., the cumulative sum of movie-field durations within a single cell, was remarkably conserved across neurons within and across brain regions (Figure 2f). Unlike movie-field durations, which differed by more than ten-fold between hippocampal and visual regions, cumulative durations were quite comparable, ranging from 6.2s (V1) to 10.2s (CA3) (Figure 2f, LGN=8.8±0.21sec, V1=6.2±0.09, AM-PM=7.8±0.09, DG=9.4±0.26, CA3=10.2±0.46, CA1=9.1±0.12, SUB=9.5±0.27). Thus, hippocampal movie-fields are longer and less multi-peaked than visual areas, such that the total duration of elevated activity was similar across all areas, spanning about a fourth of the movie, comparable to the fraction of large environments in which place cells are active^61,63,64^. To quantify the net activity in the movie-fields, we computed the total firing in the movie-fields (i.e., the area under the curve for the duration of the movie-fields), normalized by the expected discharge from the shuffled response. Unlike the ten-fold variation of movie-field durations, net movie-field discharge was more comparable (<3x variation) across brain areas, but maximal in V1 and least in subiculum (Figure 2g).

Many movie-fields showed elevated activity spanning up to several seconds, suggesting rate-code like encoding (Figure 2h). However, some cells showed movie-fields with elevated spiking restricted to less than 50ms, similar to responses to briefly flashed stimuli in anesthetized cats^12,13,66^. This is suggestive of a temporal code, characterized by low spike timing jitter^67^. Such short-duration movie-fields were not only common in the thalamus (LGN), but also AM-PM, three synapses away from the retina. A small fraction of cells in the hippocampal areas, more than five synapses away from the retina, showed such temporally coded fields as well (Figure 2h).

To determine the stability and temporal-continuity of movie tuning across the neural ensembles we computed the population vector overlap between even and odd trials^68^ (see *Methods*). Population response stability was significantly greater for tuned than for untuned neurons (Figure 3-figure supplement 1). The population vector overlap around the diagonal was broader in hippocampal regions than visual cortical and LGN, indicating longer temporal-continuity, reflective of their longer movie-fields. Further, the population vector overlap away from the diagonal was larger around frames 400-800 in all brain areas due to the longer movie-fields in that movie segment (see below).

**Figure 3.**
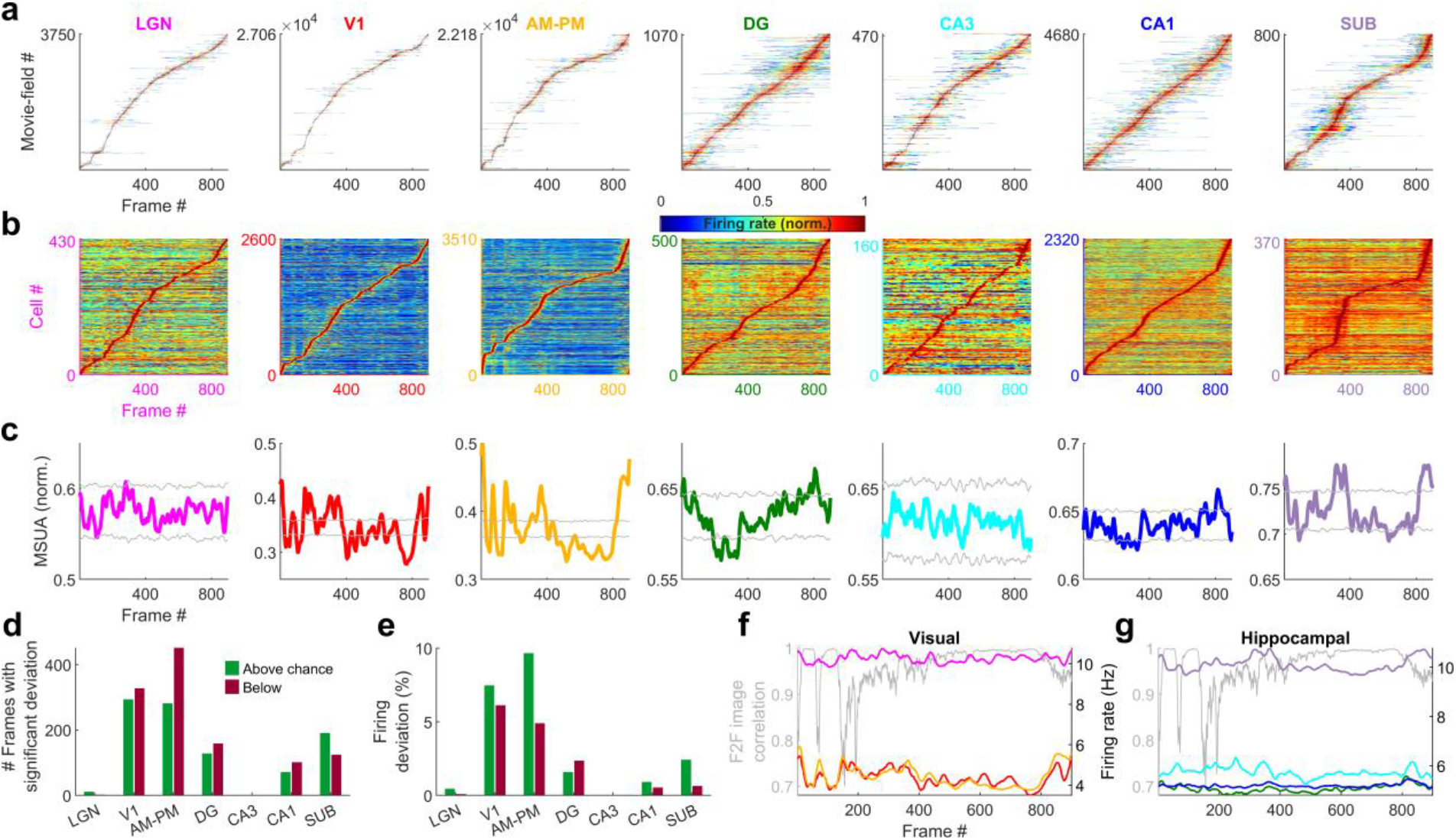
| Population averaged movie-tuning varies across brain areas. **(a)** Stack plot of all the movie-fields detected from all tuned neurons of a brain region. Color indicates relative firing rate, normalized by the maximum firing rate in that movie-field. The movie-fields were sorted according to the frame with the maximal response. Note accumulation of fields in certain parts of the movie, especially in subiculum and AM-PM. **(b)** Similar to (a), but using only a single, tallest movie-field peak from each neuron showing a similar pattern, with pronounced overrepresentation of some portions of the movie in most brain areas. Each neuron’s response was normalized by its maximum firing rate. The average firing rate of non-peak frames, which was inversely related to the depth of modulation, was smallest (0.35*x* of the average peak response across all neurons) for V1, followed by AM-PM 0.37, leading to blue shades. Average non-peak responses were higher for other regions (0.57x the peak for LGN, CA3-0.61, DG-0.62, CA1-0.64 and SUB-0.76), leading to warmer off-diagonal colors. **(c)** Multiple single unit activity (MSUA) in a given brain region, obtained as the average response across all tuned cells, by using maxima-normalized response for each cell from (b). Gray lines indicate mean±4*std response from the shuffle data corresponding to *p*=0.025 after Bonferroni correction for multiple comparisons (see *Methods*). AM-PM had the largest MSUA modulation (sparsity=0.01) and CA1 had the smallest (sparsity=1.8×10^-4^). The MSUA modulation across several brain region pairs –AM&PM-DG, V1-CA3, DG-CA3, CA3-CA1 and CA1-SUB were not significantly correlated (Pearson correlation coefficient *p*>0.05). Some brain region pairs, DG-LGN, DG-V1, AM&PM-CA3, LGN-CA1, V1-CA1, DG-SUB and CA3-SUB, were significantly negatively correlated (*r*<-0.18, *p*<4.0×10^-7^). All other brain region pairs were significantly positively correlated (*r*>0.07, *p*<0.03). **(d)** Number of frames for which the observed MSUA deviates from the z=±4 range from (c), termed significant deviation. V1 and AM-PM had the largest positive deviant frames (289), and CA3 had the least (zero). Unlike CA3, the low number of deviant frames for LGN could not be explained by sample size, because there were more tuned cells in LGN than SUB. **(e)** Firing in deviant frames above (or below) chance level, as a percentage of the average response. Above chance level deviation was greater or equal to that below, for all brain regions, with the largest positive deviation in AM-PM (9.3%), largest negative deviation in V1 (6.0%), and least in CA3 (zero each). **(f)** Total firing rate response of visual regions across tuned neurons. All regions had significant negative correlation (*r*<-0.39, *p*<3.4×10^-34^) between the ensemble response and the frame-to-frame (F2F) image correlation (gray line, y-axis on the left) across movie frames. **(g)** Similar to (f), for hippocampal regions. CA3 response were not significantly correlated with the frame-to-frame correlation, dentate gyrus (*r*=0.26, *p*=4.0×10^-15^) and CA1 (*r*=0.21, *p*=1.5×10^-10^) responses were positively correlated, and subiculum response was negatively correlated (*r*=-0.44, *p*=2.2×10^-43^). Note the substantially higher mean firing rates of LGN in (f) and subiculum neurons in (g) (colored lines closer to the top) compared to other brain areas.

### Relationship between movie image content and neural movie tuning

Are all movie frames represented equally by all brain areas? The duration and density of movie-fields varied as a function of the movie frame and brain region (Figure 3-figure supplement 2). We hypothesized that this variation could correspond to the change in visual content from one frame to the next. Hence, we quantified the similarity between adjacent movie frames as the correlation coefficient between corresponding pixels and termed it as frame-to-frame (F2F) image correlation. For comparison, we also quantified the similarity between the neural responses to adjacent frames (F2F neural correlation), as the correlation coefficient between the firing rate response of neuronal ensembles between adjacent frames. For all brain regions, the neural F2F was correlated with image F2F, but this correlation was weaker in hippocampal output regions (CA1 and SUB) than visual regions like LGN and V1. The majority of brain regions had substantially reduced density of movie-fields between the movie frames 400 to 800, but the movie-fields were longer in this region. This effect as well was greater in the visual regions than hippocampal regions. Using significantly tuned neurons, we computed the average neural activity in each brain region at each point in the movie (see *Methods*). Although movie-fields (Figure 3a), or just the strongest movie-field per cell (Figure 3b), covered the entire movie, the peak normalized, ensemble activity level of all brain regions showed significant overrepresentation, i.e., deviation from the uniformity, in certain parts of the movie (Figure 3c, see *Methods*). This was most pronounced in V1 and the higher visual areas AM-PM. The number of movie frames with elevated ensemble activity was higher in visual cortical areas than hippocampal regions (Figure 3d), and also this modulation (see *Methods*) was smaller in hippocampus and LGN, compared to the visual cortical regions (Figure 3e).

Using the significantly tuned neurons, we also computed the average neural activity in each brain region corresponding to each frame in the movie, without peak rate normalization (see *Methods*). The degree of continuity between the movie frames, quantified as above (F2F image correlation), was inversely correlated with the ensemble rate modulation in all areas except DG, CA3 and CA1 (Figure 3f and g). As expected for a continuous movie, this F2F image correlation was close to unity for most frames, but highest in the latter part of the movie where the images changed more slowly. The population wide elevated firing rates, as well as the smallest movie-fields, occurred during the earlier parts (Figure 3-figure supplement 2). Thus, the movie-code was stronger in the segments with greatest change across movie frames, in agreement with recent reports of visual cortical encoding of flow stimuli^69^. These results show differential population representation of the movie across brain regions.

### Differential neural encoding of sequential versus scrambled movie in visual and hippocampal areas

If these responses were purely visual, a movie made of scrambled sequence of images would generate equally strong or even stronger selectivity due to the even larger change across movie frames, despite the absence of similarity between adjacent frames. To explore this possibility, we investigated neural selectivity when the same movie frames were presented in a fixed but scrambled sequence (scrambled movie, Figure 4-Video 1). The within frame and the total visual content was identical between the continuous and scrambled movies, and the same sequence of images was repeated many times in both experiments (see *Methods*). But there was no correlation between adjacent frames, i.e., visual continuity, in the latter (Figure 4a).

**Figure 4.**
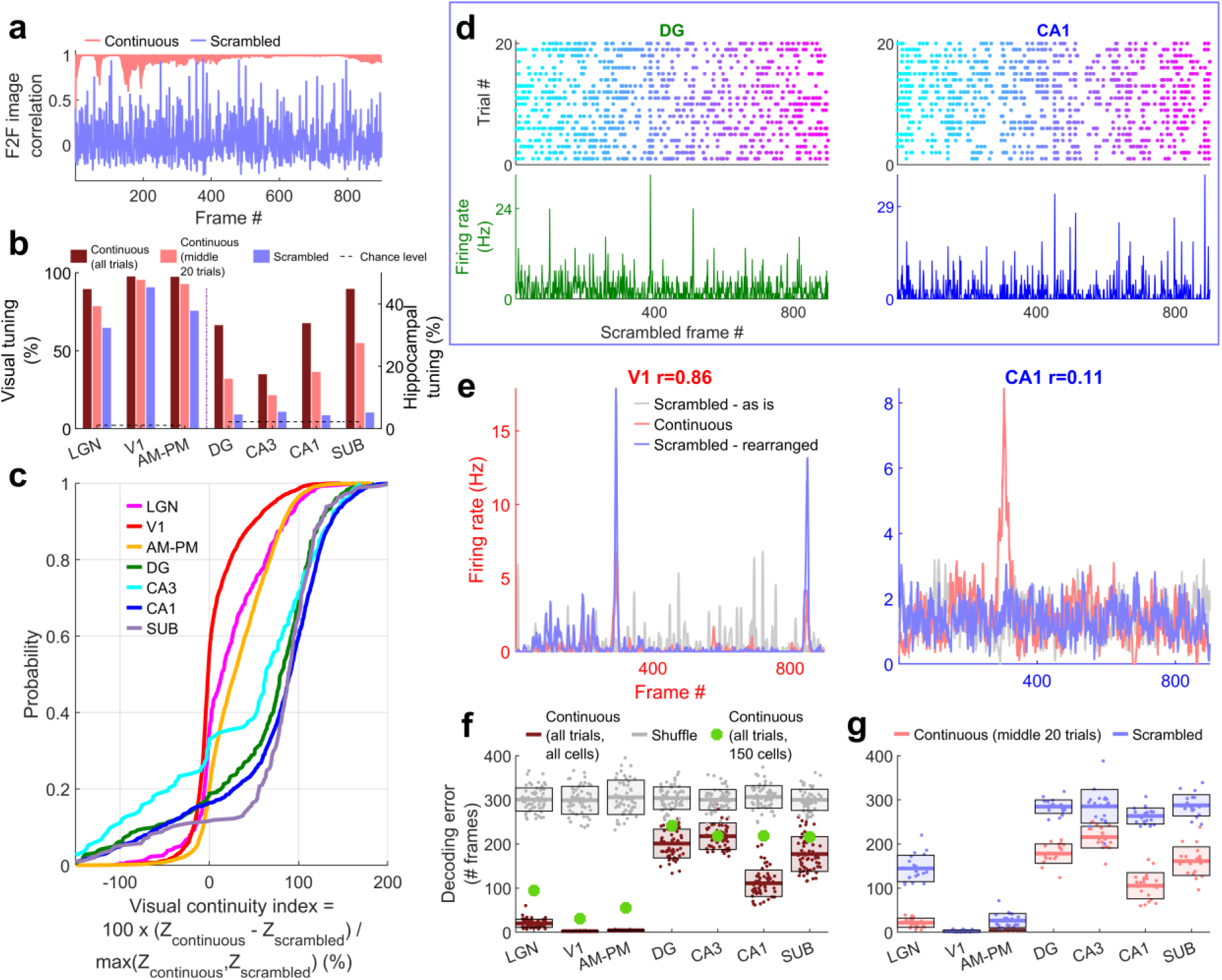
| Larger reduction of selectivity in hippocampal than visual regions due to scrambled presentation. **(a)** Similarity between the visual content of one frame with the subsequent one, quantified as the *Pearson* correlation coefficient between pixel-pixel across adjacent frames for the continuous movie (pink) and the scrambled sequence (lavender), termed F2F image correlation. Similar to Figure 3g. For the scrambled movie, the frame number here corresponded to the chronological frame sequence, as presented. **(b)** Fraction of broad spiking neurons significantly modulated by the continuous movie (red) or the scrambled sequence (blue) using z-scored sparsity measures (similar to Figure 1, see *Methods*). For all brain regions, continuous movie generated greater selectivity than scrambled sequence (KS-test *p*<7.4×10^-4^). **(c)** Percentage change in the magnitude of tuning between the continuous and scrambled movies for cells significantly modulated by either continuous or scrambled movie, termed visual continuity index. The largest drop in selectivity due to scrambled movie occurred in CA1 (90.3±2.0%), and least in V1 (−1.5±0.6%). Visual continuity index was significantly different between all brain region pairs (KS-test *p*<0.03) and significantly greater for hippocampal areas than visual (8.2-fold, *p*<10^-100^). **(d)** Raster plots (top) and mean rate responses (color, bottom) showing increased spiking responses to only one or two scrambled movie frames, lasting about 50ms. Tuned responses to scrambled movie were found in all brain regions, but these were the least frequent in DG and CA1. **(e)** One representative cell each from V1 (left) and CA1 (right), where the frame rearrangement of scrambled responses resulted in a response with high correlation to the continuous movie response for V1, but not CA1. *Pearson* correlation coefficient values of continuous movie and rearranged scrambled responses are indicated on top. **(f)** Average decoding error for observed data (see *Methods*), over 60 trials for continuous movie (maroon), was significantly lower than shuffled data (gray) (KS-test *p*<1.2×10^-22^). Solid line – mean error across 60 trials using all tuned cells from a brain region, shaded box – s.e.m., green dots – mean error across all trials using a random subsample of 150 cells from each brain region. Decoding error was lowest for V1 (30.9 frames) and highest in DG (241.2) and significantly different between all brain regions pairs (*p*<1.9×10^-4^), except CA3-CA1, CA3-subiculum and CA1-subiculum (*p*>0.63). **(g)** Similar to (f), decoding of scrambled movie was significantly worse than that for the continuous movie (KS-test *p*<2.6×10^-3^). Scrambled responses, in their “as is”, chronological order were used herein. LGN decoding error for scrambled presentation was 6.5x greater than that for continuous movie, whereas the difference in errors was least for V1 (1.04x). Scrambled movie decoding error for all visual areas and for CA1 and subiculum was significantly smaller than chance level (KS-test *p*<2.6×10^-3^), but not DG and CA3 (*p*>0.13). Only the middle 20 trials of the continuous movie were used for comparison with the scrambled movie since the scrambled movie was only presented 20 times. Middle trials of the continuous movie were chosen as the appropriate subset since they were chronologically closest to the scrambled movie presentation.

For all brain regions investigated, the continuous movie generated significantly greater modulation of neural activity than the scrambled sequence (Figure 4b). Middle 20 trials of the continuous movie were chosen as the appropriate subset for comparison since they were chronologically closest to the scrambled movie presentation. This choice ensured that other long-term effects, such as behavioral state change, instability of single unit measurement and representational^70^ or behavioral^71^ drift could not account for the differences in neural responses to continuous and scrambled movie presentation. This preference for continuous over scrambled movie was the greatest in hippocampal regions where the percentage of significantly tuned neurons (4.4%, near chance level of 2.3%) reduced more than 4-fold compared to the continuous movie (17.8%, after accounting for the lesser number of trials, see *Methods*). This was unlike visual areas where the scrambled (80.4%) and the continuous movie (92.4%) generated similar prevalence levels of selectivity (Figure 4b). The few hippocampal cells which had significant selectivity to the scrambled sequence, did not have long-duration responses, but only very short, ∼50ms long responses (Figure 4d), reminiscent of, but even sharper than human hippocampal responses to flashed images^33^. To estimate the effect of continuous movie compared to the scrambled sequence on individual cells, we computed the normalized difference between the continuous and scrambled movie selectivity for cells which were selective in either condition (Figure 4c, see *Methods*). This visual continuity index was more than eight-fold higher in hippocampal areas (median values across all 4 hippocampal regions = 87.8%) compared to the visual areas (median = 10.6% across visual regions).

The pattern of increasing visual continuity index as we moved up the visual hierarchy, largely paralleled the anatomic organization^72^, with the greatest sensitivity to visual continuity in the hippocampal output regions, CA1 and subiculum, but there were notable exceptions. The primary visual cortical neurons showed the least reduction in selectivity due to the loss of temporally contiguous content, whereas LGN neurons, the primary source of input to the visual cortex and closer to the periphery, showed far greater sensitivity (Figure 4c).

Many visual cortical neurons were significantly modulated by the scrambled sequence, but their number of movie-fields per cell was greater and their duration was shorter than during the continuous movie (Figure 4-figure supplement 1&2). This could occur due to the loss of frame-to-frame correlation in the scrambled sequence. The average activity of the neural population in V1 and AM-PM showed significant deviation even with the scrambled movie, comparable to the continuous movie, but this multi-unit ensemble response was uncorrelated with the frame-to-frame correlation in the scrambled sequence (Figure 4-figure supplement 3). A substantial fraction of visual cortical and LGN responses to the scrambled sequence could be rearranged to resemble continuous movie responses (Figure 4-figure supplement 4, see *Methods*). The latency needed to shift the responses was least in LGN and largest in AM-PM, as expected from the feed-forward anatomy of visual information processing^36,72^ (Figure 4-figure supplement 4). Unlike visual areas, such rearrangement did not resemble the continuous movie responses in the hippocampal regions (example cells in Figure 4e, also see Figure 4-figure supplement 4 for statistics and details). Further, even after rearranging the hippocampal responses, their selectivity to the scrambled movie presentation remained near chance levels (Figure 4-figure supplement 5).

Population vector decoding of the ensemble of a few hundred place cells is sufficient to decode the rat’s position using place cells^73^, and the position of a passively moving object^40^. Using similar methods, we decoded the movie frame number (see *Methods*). Continuous movie decoding was better than chance in all brain regions analyzed (Figure 4f). Upon accounting for the number of tuned neurons from different brain regions, the decoding was most accurate in V1, and least in dentate gyrus. Scrambled movie decoding was significantly weaker yet above chance level (based on shuffles, see *Methods*) in visual areas, but not in CA3 and dentate gyrus. But CA1 and subiculum neuronal ensembles could be used to decode scrambled movie frame number slightly above chance levels (Figure 4g). Similarly, the population overlap between even and odd trials for the scrambled sequence was strong for visual areas, and weaker in hippocampal regions, but significantly greater than untuned neurons in hippocampal regions (Figure 4-figure supplement 6). Combined with the handful of neurons in hippocampus whose movie selectivity persisted to the scrambled presentation, this suggests that loss of correlations between adjacent frames in the scrambled sequence abolishes most, but not all of the hippocampal selectivity to visual sequences.

## Discussion

### Movie tuning in the visual areas

To understand how neurons encode a continuously unfolding visual episode, we investigated the neural responses in the head fixed mouse brain to an isoluminant, black-and-white, silent human movie, without any task demands or rewards. As expected, neural activity showed significant modulation in all thalamo-cortical visual areas, with elevated activity in response to specific parts of the movie, termed movie-fields. Most (96.6%, 6554/6785) of thalamo-cortical neurons showed significant movie tuning. This is nearly double that reported for the classic stimuli such as Gabor patches in the same dataset^36^, although a direct comparison is difficult due to the differences in experimental and analysis methods. For example, the classic stimuli were presented for 250ms, preceded by a blank background whereas the images changed every 30ms in a movie. On the other hand, significant tuning of the vast majority of visual neurons to movies is consistent with other reports^11–13,15,17,66,69–71^. Thus, movies are a reliable method to probe the function of the visual brain and its role in cognition.

### Movie tuning in hippocampal areas

Remarkably, a third of hippocampal neurons (32.9%, 3379/10263) were also movie-tuned, comparable to the fraction of neurons with significant spatial selectivity in mice^74^ and bats^75^, and far greater than significant place cells in the primate hippocampus^76–78^. While the hippocampus is implicated in episodic memory, rodent hippocampal responses are largely studied in the context of spatial maps or place cells, and more recently in other tasks which requires active locomotion or active engagement^7,79^. However, unlike place cells^5,6^, movie-tuning remained intact during immobility in all brain areas studied, which could be because self-motion causes consistent changes in multisensory cues during spatial exploration but not during movie presentation. This dissociation of the effect of mobility on spatial and movie selectivity agrees with the recent reports of dissociated mechanisms of episodic encoding and spatial navigation in human amnesia^80^. Our results are broadly consistent with prior studies that found movie selectivity in human hippocampal single neurons^81^. However, that study relied on famous, very familiar movie clips, similar to the highly familiar image selectivity^33^ to probe episodic memory recall. In contrast, mice in our study had seen this black-and-white, human movie clip only in two prior habituation sessions and it is very unlikely that they understood the episodic content of the movie. Recent studies found human hippocampal activation in response to abrupt changes between different movie clips^34,82,83^, which is broadly consistent with our findings. Future studies can investigate the nature of hippocampal activation in mice in response to familiar movies to probe episodic memory and recall. These observations support the hypothesis that specific visual cues can create reliable representations in all parts of hippocampus in rodents^5,37,40^, nonhuman primates^76,78^ and humans^84,85^, unlike spatial selectivity which requires consistent information from multisensory cues^28,38,86^.

### Mega-scale nature of movie-fields

Across all brain regions, neurons showed a mega-scale encoding by movie-fields varying in duration by up to 1000-fold, similar to, but far greater than recent reports of 10-fold multi-scale responses in the hippocampus^59–64,87^. While neural selectivity to movies has been studied in visual areas, such mega-scale coding has not been reported. Remarkably, mega-scale movie coding was found not only across the population but even individual LGN and V1 neurons could show two different movie fields, one lasting less than 100ms and other exceeding 10,000ms. The speed at which visual content changed across movie frames could explain a part, but not all of this effect. The mechanisms governing the mega-scale encoding would require additional studies. For example, the average duration of the movie-field increased along the feed-forward hierarchy, consistent with the hierarchy of response lags during language processing^88^. Paradoxically, the mega-scale coding of movie field meant the opposite pattern also existed, with 10s long movie fields in some LGN cells while less than 100ms long movie fields in subiculum.

### Continuous versus scrambled movie responses

The analysis of scrambled movie-sequence allowed us to compute the neural response latency to movie frames. This was highest in AM-PM (91ms) than V1 (74ms) and least in LGN (60ms), thus following the visual hierarchy. The pattern of movie tuning properties was also broadly consistent between V1 and AM/PM (Fig 2). However, several aspects of movie-tuning did not follow the feed-forward anatomical hierarchy. For example, all metrics of movie selectivity (Fig 2) to the continuous movie showed a consistent pattern that was the inconsistent to the feed-forward anatomical hierarchy: V1 had stronger movie tuning, higher number of movie fields per cell, narrower movie-field widths, larger mega-scale structure, and better decoding than LGN. V1 was also more robust to scrambled sequence than LGN. One possible explanation is that there are other sources of inputs to V1, beyond LGN, that contribute significantly to movie tuning^89^. Amongst the hippocampal regions, the tuning properties of CA3 neurons (field durations, mega-chronicity index, visual continuity index and several measures of population modulation) were closest to that of visual regions, even though the prevalence of tuning in CA3 was lesser than that in other hippocampal as well as visual areas.

### Emergence of episode-like movie code in hippocampus

Temporal integration window^90–92^ as well as intrinsic timescale of firing^36^ increase along the anatomical hierarchy in the cortex, with the hippocampus being farthest removed from the retina^72^. This hierarchical anatomical organization, with visual areas being upstream of hippocampus could explain the longer movie-fields, the strength of tuning, number of movie peaks, their width and decoding accuracy in hippocampal regions. This could also explain the several fold greater preference for the continuous movie over scrambled sequence in the hippocampus compared to the upstream visual areas. But, unlike reports of image-association memory in the inferior temporal cortex for unrelated images^93,94^, only a handful hippocampal neurons showed selective responses to the scrambled sequence. These results, along with the longer duration of hippocampal movie-fields could mediate visual-chunking or binding of a sequence of events. In fact, evidence for episodic-like chunking of visual information was found in all visual areas as well, where the scrambled-sequence not only reduced neural selectivity but caused fragmentation of movie-fields (Figure 4-figure supplement 4).

### No evidence of nonspecific effects

Could the brain-wide mega-scale tuning be an artifact of poor unit isolation, e.g., due to an erroneous mixing of two neurons, one with very short and another with very long movie-fields? This is unlikely since the LGN and visual cortical neural selectivity to classic stimuli (Gabor patches, drifting gratings etc.) in the same dataset was similar to that reported in most studies^36^ whereas poor unit isolation should reduce these selective responses. However, to directly test this possibility, we calculated the correlation between the unit isolation index (or fraction of refractory violations) and the mega-scale index of the cell, while factoring out the contribution of mean firing rate (Figure 1-figure supplement 8). This correlation was not significant (*p*>0.05) for any brain areas.

### Movie-fields vs. place-fields

Do the movie fields arise from the same mechanism as place fields? Studies have shown that when rodents are passively moved along a linear track that they had explored^6^, or when the images of the environment around a linear track was played back to them^5^, some hippocampal neurons generated spatially selective activity. Since the movie clip involved change of spatial view, one could hypothesize that the movie fields are just place fields generated by passive viewing. This is unlikely for several reasons. Mega-scale movie fields were found in the vast majority of all visual areas including LGN, far greater than spatially modulated neurons in the visual cortex during virtual navigation^95,96^. Further, in prior passive viewing experiments, the rodents were shown the same narrow linear track, like a tunnel, that they had previously explored actively to get food rewards at specific places. In contrast, in current experiments, these mice had never actively explored the space shown in the movie, nor obtained any rewards. Active exploration of a maze, combined with spatially localized rewards engages multisensory mechanisms resulting in increased place cell activation^22,28,97^ which are entirely missing in these experiments during passive viewing of a movie, presented monocularly, without any other multisensory stimuli and without any rewards. Compared to spontaneous activity about half of CA1 and CA3 neurons shutdown during spatial exploration and this shutdown is even greater in the dentate gyrus. Further, compared to the exploration of a real-world maze, exploration of a visually identical virtual world causes 60% reduction in CA1 place cell activation^86^. In contrast, there was no evidence of neural shutdown during the movie presentation compared to grey screen spontaneous epochs (Figure 1-figure supplement 8). Similarly, the number of place fields (in CA1) per cell on a long track is positively correlated with the mean firing rate of the cell^62^, which was not seen here for CA1 movie fields.

A recent study showed that CA1 neurons encode the distance, angle, and movement direction of motion of a vertical bar of light^40^, consistent with the position of hippocampus in the visual circuitry^72^. Do those findings predict the movie tuning herein? There are indeed some similarities between the two experimental protocols –purely passive optical motion without any self-motion or rewards. However, there are significant differences too; similar to place cells in the real and virtual worlds^38^, all the cells tuned to the moving bar of light had single receptive fields with elevated responses lasting a few seconds; there were neither punctate responses nor even 10-fold variation in neural field durations, let alone the 1000-fold change reported here. Finally, those results were reported only in area CA1, while the results presented here cover nearly all the major stations of the visual hierarchy.

Notably, hippocampal neurons did not encode Gabor patches or drifting gratings in the same dataset, indicating the importance of temporally continuous sequences of images for hippocampal activation^36^. This is consistent with the hypothesis that the hippocampus is involved in coding spatial sequences^21,29,98^. However, unlike place cells that degrade in immobile rats, hippocampal movie tuning was unchanged in the immobile mouse. Further, the scrambled sequence too was presented in the same sequence many times, yet movie tuning dropped to chance level in the hippocampal areas. Unlike visual areas, scrambled sequence response of hippocampal neurons could not be rearranged to obtain the continuous movie response. This shows the importance of continuous, episodic content instead of mere sequential recurrence of unrelated content for rodent hippocampal activation. We hypothesize that similar to place cells, movie-field responses without task-demand would play a role, to be determined, in episodic memory. Further work involving a behavior report for the episodic content can potentially differentiate between the sequence coding described here and the contribution of episodically meaningful content. However, the nature of movie selectivity tested so far in humans was different (recall of famous, short movie clips^81^, or at event boundaries^34^) than in rodents here (human movie, selectivity to specific movie segments).

### Broader outlook

Our findings open up the possibility of studying thalamic, cortical, and hippocampal brain regions in a simple, passive, and purely visual experimental paradigm and extend comparable convolutional neural networks^11^ to have the hippocampus at the apex^72^. Further, our results here bridge the long-standing gap between the hippocampal rodent and human studies^34,99–101^, where natural movies can be decoded from fMRI signals in immobile humans^102^. This brain-wide mega-scale encoding of a human movie episode and enhanced preference for visual continuity in the hippocampus compared to visual areas supports the hypothesis that the rodent hippocampus is involved in non-spatial episodic memories, consistent with classic findings in humans^1^ and in agreement with a more generalized, representational framework^103,104^ of episodic memory where it encodes temporal patterns. Similar responses are likely across different species, including primates. Thus, movie-coding can provide a unified platform to investigate the neural mechanisms of episodic coding, learning and memory.

## Supporting information

Episodic movie

Scrambled movie sequence

## Methods

### Experiments

We used the Allen Brain Observatory – Neuropixels Visual Coding dataset (© 2019 Allen Institute, https://portal.brain-map.org/explore/circuits/visual-coding-neuropixels). This website and related publication^36^ contain detailed experimental protocol, neural recording techniques, spike sorting etc. Data from 24 mice (16 males, n=13-C57BL/6J wild-type, n=2 Pvalb-IRES-Cre×Ai32, n=6 Sst-IRES-Cre×Ai32, and n=3 Vip-IRES-Cre×Ai32) from the “Functional connectivity” dataset was analyzed herein. Prior to implantation with Neuropixel probes, mice passively viewed the entire range of images including drifting gratings, Gabor patches and movies of interest here. Videos of the body and eye movements were obtained at 30Hz and synced to the neural data and stimulus presentation using a photodiode. Movies were presented monocularly on an LCD monitor with a refresh rate of 60Hz, positioned 15cm away from the mouse’s right eye and spanned 120°x95°. 30 trials of the continuous movie presentation were followed by 10 trials of the scrambled movie. Next was a presentation of drifting gratings, followed by a quiet period of 30 minutes where the screen was blank. Then the second block of drifting gratings, scrambled movie and continuous movie was presented. After surgery, all mice were single-housed and maintained on a reverse 12-h light cycle in a shared facility with room temperatures between 20 and 22 °C and humidity between 30 and 70%. All experiments were performed during the dark cycle.

Neural spiking data was sampled at 30 kHz with a 500Hz high pass filter. Spike sorting was automated using Kilosort2^105^. Output of Kilosort2 was post-processed to remove noise units, characterized by unphysiological waveforms. Neuropixel probes were registered to a common co-ordinate framework^106^. Each recorded unit was assigned to a recording channel corresponding to the maximum spike amplitude and then to the corresponding brain region. Broad spiking units identified as those with average spike waveform duration (peak to trough) between 0.45 to 1.5ms and those with mean firing rates above 0.5Hz were analyzed throughout, except Figure 1-figure supplement 8.

### Movie tuning quantification

The movie consisted of 900 frames: 30s total, 30Hz refresh rate, 33.3ms per frame. At the first level of analysis, spike data were split into 900 bins, each 33.3ms wide (the bin size was later varied systematically to detect mega-scale tuning, see below). The resulting tuning curves were smoothed with a Gaussian window of σ=66.6 ms or 2 frames. The degree of modulation and its significance was estimated by the sparsity *s* as below, and as previously described^40,86^.

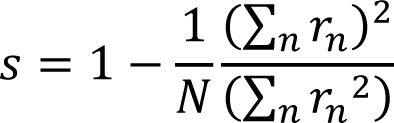

where *r_n_* is the firing rate in the 𝑛𝑡ℎ frame or bin and N=900 is the total number of bins. This is equivalent to “lifetime sparseness”, used previously^11,14^, except for the normalization factor of (1-1/N), which is close to unity, when N is close to 900 as in the case of movies. Statistical significance of sparsity was computed using a bootstrapping procedure, which does not assume a normal distribution. Briefly, for each cell, the spike train as a function of the frame number from each trial was circularly shifted by different amounts and the sparsity of the randomized data computed. This procedure was repeated 100 times with different amounts of random shifts. The mean value and standard deviation of the sparsity of randomized data were used to compute the z-scored sparsity of observed data using the function *zscore* in MATLAB. The observed sparsity was considered statistically significant if the z-scored sparsity of the observed spike train was greater 2, which corresponds to *p*<0.023 in a one tailed t-test. A similar method was used to quantify significance of the scrambled movie tuning, as well as for the subset of data with only stationary epochs, or its equivalent subsample (see below). Middle 20 trials of the continuous movie were used in comparisons with the scrambled movie in Figure 4, to ensure a fair comparison by using same number of trials, with similar time delays across measurements.

In addition to sparsity, we quantified movie tuning using two other measures.

Depth of modulation = (𝑟_𝑚𝑎𝑥_ - 𝑟_𝑚𝑖𝑛_)/ (𝑟_𝑚𝑎𝑥_ + 𝑟_𝑚𝑖𝑛_), where 𝑟_𝑚𝑎𝑥_ and 𝑟_𝑚𝑖𝑛_ are the largest and lowest firing rates across movie frames, respectively.

Mutual information 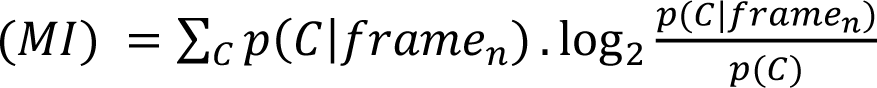

Where 𝑝(𝐶) = ∑_𝑛_ 𝑝(𝑓𝑟𝑎𝑚𝑒_𝑛_). 𝑝(𝐶|𝑓𝑟𝑎𝑚𝑒_𝑛_) and 𝐶 is the average spike count in 0.033 second window which corresponds to 1 movie frame. 𝑝(𝑓𝑟𝑎𝑚𝑒_𝑛_) is 1/900, as all frames were presented equal number of times. Statistical significance of these alternative measures of selectivity was computed similar to that for sparsity and is detailed in Figure 1-figure supplement 3.

### Stationary epoch and sharp wave ripple free epoch identification

To eliminate the confounding effects of changes in behavioral state associated with running, we repeated our analysis in stationary epochs, defined as epochs when the running speed remained less than 2cm/sec for this period, as well as for at least 5 seconds before and after this period. Analysis was further restricted to sessions with at least 5 total minutes of these epochs during the 60 trials of continuous movie presentation. To account for using lesser data of the stationary epochs, we compared the tuning using a random subsample of data, regardless of running or stopping and compared the two results for difference in selectivity.

Similarly, to remove epochs of sharp wave ripples (SWR), we first computed band passed power in the hippocampal (CA1) recording sites in the 150-250Hz range. SWR occurrence was noted if any of the best 5 sites in CA1 (those with highest theta (5-12Hz) to delta (1-4Hz) ratio), or the median SWR across all CA1 sites exceeded their respective 3 standard deviations of power. To remove SWRs, we removed frames corresponding to ±0.5second around the SWR occurrence and recomputed movie tuning in the remaining data. Similar to the stationary epoch calculation above, we compared tuning to an equivalent random subset to account for loss of data.

### Pupil dilation and theta power comparisons

To assess the contribution of arousal state on movie tuning, we re-calculated z-scored sparsity in epochs with high vs. low pupil dilation. The pupil was tracked at a 30Hz sampling rate, and the height and width of the elliptical fit as provided in the publicly available dataset was used. For each session, the pupil area thus calculated was split into two equal halves, by using data above and below the 50^th^ percentile. The resultant z-scored sparsity is reported in Figure 1-figure supplement 7.

Similarly, the theta power computed from the band passed local field potential signal in the 5-12Hz range was split into 2 equal data sub segments. The channel from CA1, with the highest average theta to delta (1-4Hz) power ratio was nominated as the channel to be used for these calculations. Movie tuning in data with high and low theta power thus separated is reported in Figure 1-figure supplement 7.

### Mega-scale movie-field detection in tuned neurons

For neurons with significant movie-sparsity, i.e., movie-tuned, the movie response was first recalculated at a higher resolution of 3.33ms (10 times the frame rate of 33.3ms). The *findpeaks* function in MATLAB was used to obtain peaks with *prominence* larger than 110% (1.1x) the range of firing variation obtained by chance, as determined from a sample shuffled response.

This calculation was repeated at different smoothing values (logarithmically spaced in 10 Gaussian smoothing schemes with σ ranging from 6.7ms to 3430ms), to ensure that long as well as short movie-fields were reliably detected and treated equally. For frames where overlapping peaks were found at different smoothing levels, we employed a comparative algorithm to only select the peak(s) with higher prominence score. This score was obtained as the ratio of the peak’s prominence to the range of fluctuations in the correspondingly smoothed shuffle. This procedure was conducted iteratively, in increasing order of smoothing. If a broad peak overlapped with multiple narrow ones, the sum of scores of the narrow ones was compared with the broad one. To ensure that peaks at the beginning as well as the end of the movie frames were reliably detected, we circularly wrapped the movie response, for the observed as well as shuffle data.

### Identifying frames with significant deviations in multiple single-unit activity (MSUA)

First, the average response across tuned neurons for each brain region was computed for each movie frame, after normalizing the response of each cell by the peak firing response. This average response was used as the observed “Multiple single unit activity (MSUA)” in Figure 3. To compute chance level, individual neuron responses were circularly shifted with respect to the movie frames to break the frame to firing rate association but maintain overall firing rate modulation. 100 such shuffles were used, and for each shuffle, the shuffled MSUA response was computed by averaging across neurons. Across these 100 shuffles, mean and standard deviation was obtained for all frames, and used to compute the z-score of the observed MSUA. To obtain significance at *p*=0.025 level, Bonferroni correction was applied, and the appropriate z-score (4.04) level was chosen. The number of frames in the observed MSUA above (and below) this level were further quantified in Figure 3. The firing deviation for these frames was computed as the ratio between the mean observed MSUA and the mean shuffled MSUA, reported as a percentage, for frames corresponding to z-score greater than +4 or less than -4. To obtain a total firing rate report, where each spike gets equal vote, we computed the total firing response by computing the total rate across all tuned neurons (and averaging by the number of neurons) in Figure 3 and across all neurons in Figure 3-figure supplement 2.

### Population Vector Overlap

To evaluate the properties of a population of cells, movie presentations were divided into alternate trials, yielding even and odd blocks^68^. Population vector overlap was computed between the movie responses calculated separately for these 2 blocks of trials. Population vector overlap between frames *x* of the even trials & frame *y* of the odd trials was defined as the Pearson correlation coefficient between the vectors *(R_1,x_, R _2,x_, … R _N,x_) & (R _1,y_, R _2,y_, … R _N,y_),* where *R _n,x_* is the mean firing rate response of the *n*^th^ neuron to the *x*^th^ movie frame. N is the total number of neurons used, for each brain region. This calculation was done for *x* and *y* ranging from 1 to 900, corresponding to the 900 movie frames. The same method was used for tuned and untuned neurons in continuous movie responses in Figure 3-figure supplement 1, and for scrambled sequence responses in Figure 4-figure supplement 6.

### Decoding analysis

Methods similar to those previously described were used^40,73^. For tuned cells, the 60 trials of continuous movie were each decoded using all other trials. Mean firing rate responses in the 59 trials for 900 frames were used to compute a “look-up” matrix. Each neuron’s response was normalized between 0 and 1. At each frame in the “observed” trial, the correlation coefficient was computed between the population vector response in this trial and the look-up matrix. The frame corresponding to the maximal correlation was denoted as the decoded frame. Decoding error was computed as the average of the absolute difference between actual and decoded frames, across the 900 frames of the movie. For comparison, shuffle data was generated by randomly shuffling the cell-cell pairing of the look-up matrix and “observed response”. To enable a fair comparison of decoding accuracy across brain regions, the tuned cells from each brain region were subsampled, and a random selection of 150 cells was used. A similar procedure was used for the 20 trials of the scrambled sequence, and the corresponding middle 20 trials of the continuous movie were used here for comparison.

### Rearranged scrambled movie analysis

To differentiate the effects of visual content versus visual continuity between consecutive frames, we compared the responses of the same neuron to the continuous movie and the scrambled sequence. In the scrambled movie, the same visual frames as the continuous movie were used, but they were shuffled in a pseudo random fashion. The same scrambled sequence was repeated for 20 trials. The neural response was first computed at each frame of the scrambled sequence, keeping the frames in the chronological order of presentation. Then the scrambled sequence of frames was rearranged to recreate the continuous movie and the corresponding neural responses computed. To address the latency between movie frame presentation and its evoked neural response, which can differ across brain regions and neurons, this calculation was repeated for rearranged scrambled sequences with variable delays between τ= -500 to +500 ms (i.e., -150 to +150 frames of 3.33ms resolution, in steps of 5 frames or 16.6ms). The correlation coefficient was computed between the continuous movie response and this variable delayed response at each delay as r_measured_ (τ) = corrcoef (R_continuous_, R_scramble-rearranged_(τ)). R_continuous_ is the continuous movie response, obtained at 3.33ms resolution and similarly, R_scramble-rearranged_ corresponds to the scrambled response after rearrangement, at the latency τ. The latency τ yielding the largest correlation between the continuous and rearranged scrambled movie was designated as the putative response latency for that neuron. This was used in Figure 4-figure supplement 4. The value of r_measured_(τ_max_) was bootstrapped using 100 randomly generated frame reassignments, and this was used to z-score r_measured_(τ_max_), with z-score > 2 as criterion for significance. The resultant z-score is reported in Figure 4-figure supplement 4.

The latency τ was rounded off for use with 33ms bins and used to rearrange actual as well as shuffled data to compute the strength of tuning for scrambled presentation. Z-scored sparsity was computed as described above. This was compared with the z-scored sparsity of continuous movie as well as the scrambled movie data, without the rearrangement, and shown in Figure 4-figure supplement 5.

## Acknowledgements

We thank the Allen Brain Institute for provision of the dataset, Dr. Josh Siegle for help with the dataset, Dr. Krishna Choudhary for proof-reading of the text and Dr. Massimo Scanziani for input and feedback. This work was supported by grants to M.R.M. by the National Institutes of Health NIH 1U01MH115746.

## Author contributions

C.S.P. performed the analyses with input from M.R.M. C.S.P. and M.R.M. wrote the manuscript.

## Data availability

All data is publicly available at the Allen Brain Observatory – Neuropixels Visual Coding dataset (© 2019 Allen Institute, https://portal.brain-map.org/explore/circuits/visual-coding-neuropixels).

## Code availability

All analyses were performed using custom-written code in MATLAB version R2020a. Codes written for analysis and visualization are available on GitHub, at https://github.com/cspurandare/ELife_MovieTuning^107^.

## Video Legends

**Figure 1-video 1 | Sequential movie.** The 30 second movie clip shown, along with the frame number indicated in the top right corner (updated every second, or every 30 frames). The same movie clip was shown in two blocks of 30 repeats each.

**Figure 4-video 1 | Scrambled movie.** Frames from the sequential video clip (Figure 1-video 1) were presented in a scrambled sequence, with the same sequence repeated 20 times (2 blocks of 10 trials each). Frame numbers in the scrambled sequence are indicated in the top right corner.

## Supplementary Figures

**Figure 1-figure supplement 1.**
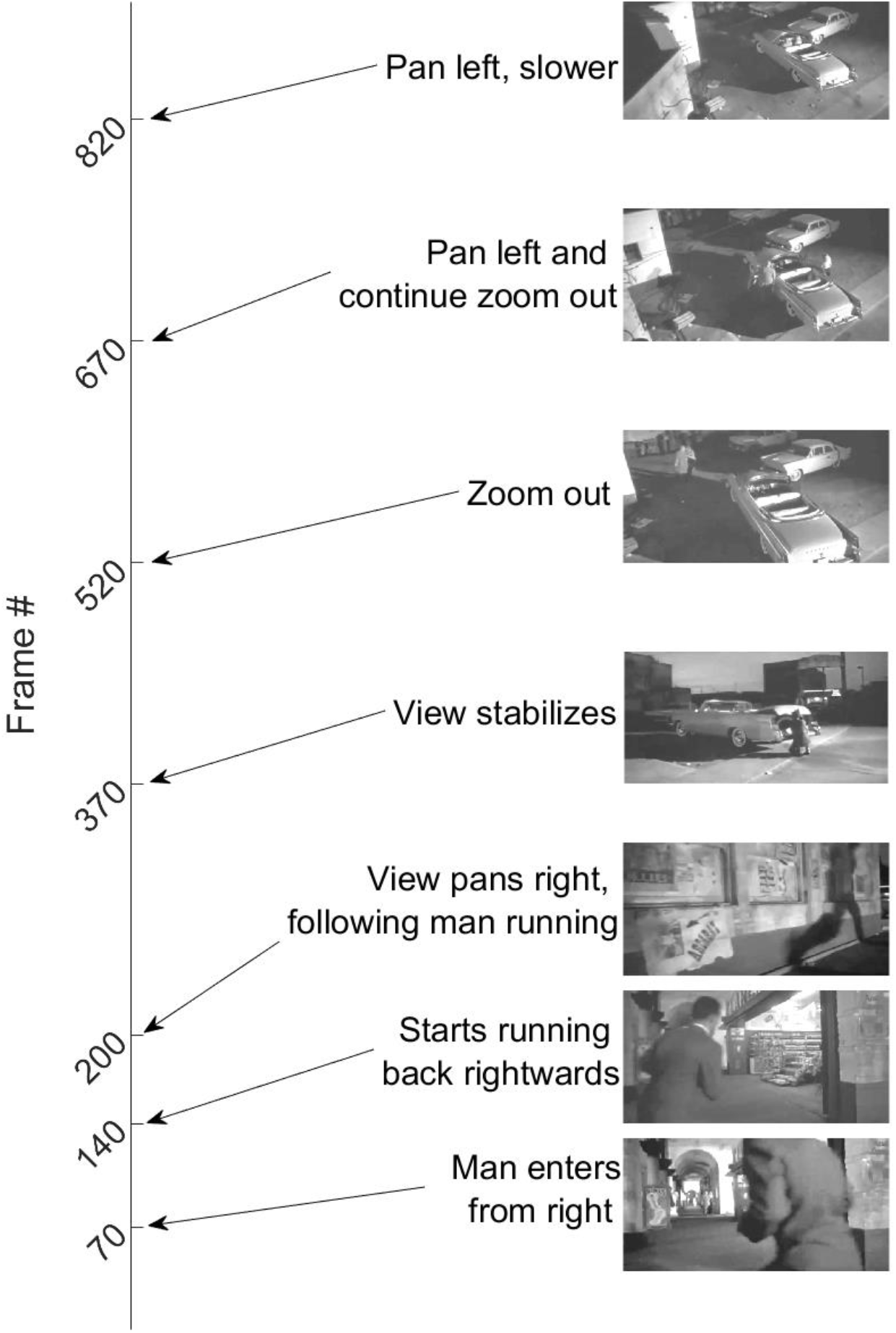
| The movie. The 30 second long, silent, black-and-white, isoluminant movie with frame numbers denoting key episodes in this continuous segment.

**Figure 1-figure supplement 2.**
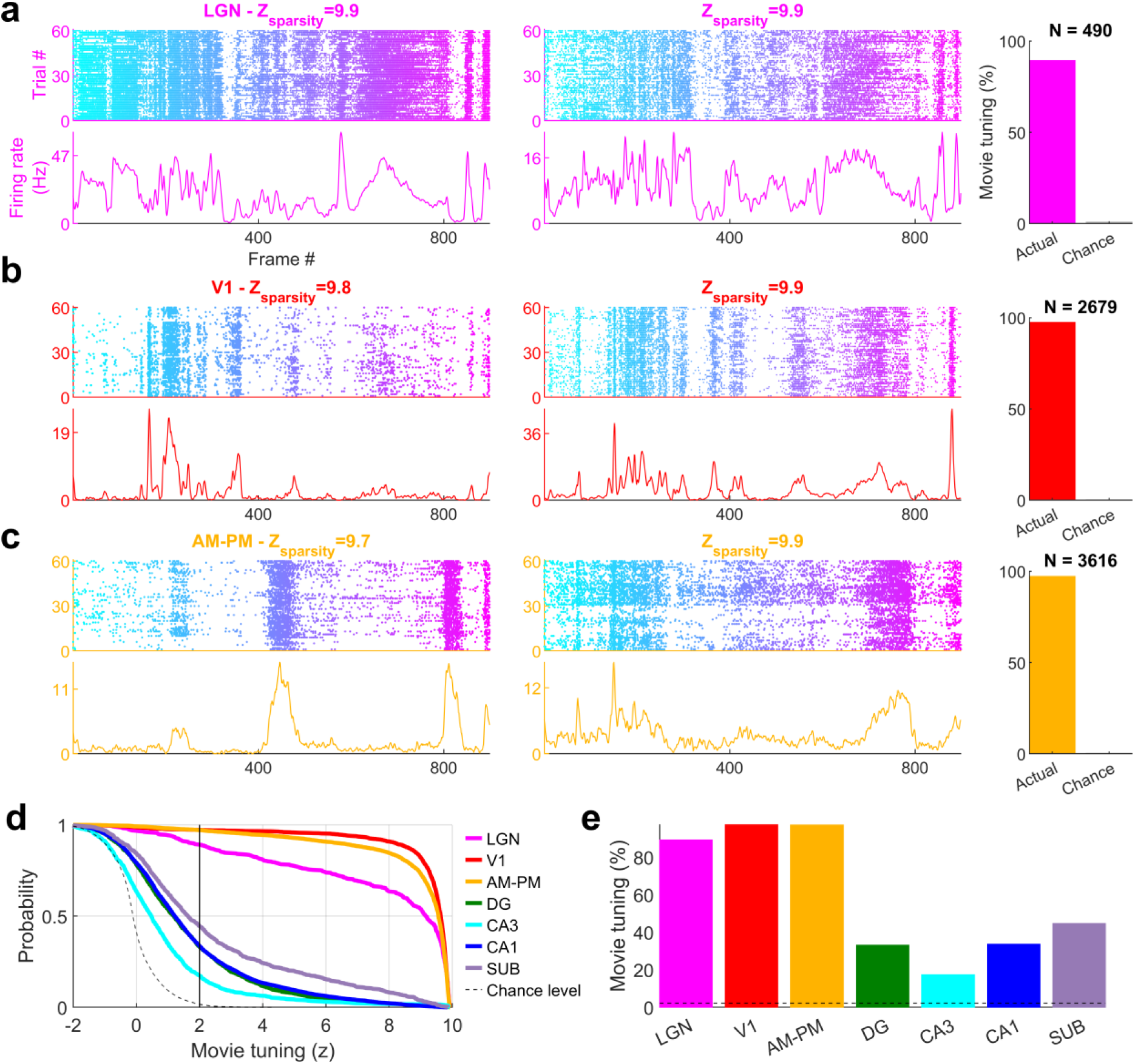
| Movie selectivity across brain areas. (a) Similar to Figure 1, representative single cells from LGN showing selective movie responses. Strength of modulation, quantified by z-scored sparsity is indicated above. The total number of broad spiking cells used (N) and the fraction selective are shown by the bar chart on the right. **(b)** Same as that (a), for V1 and **(c)** for higher visual areas AM-PM. **(d)** Cumulative distribution of movie selectivity across all broad spiking cells, including significantly (z>2 vertical black line, see *Methods*) tuned cells. The largest prevalence of selectivity in broad spiking neurons was seen in primary visual cortex (V1, 97.3%, 2606 out of 2679) and least in CA3 hippocampus (17.3%, 168 out of 969). **(e)** All brain regions analyzed showed far greater selectivity than the chance level (dashed gray line). There was a clear difference in the strength of movie-tuning between visual and hippocampal areas.

**Figure 1-figure supplement 3.**
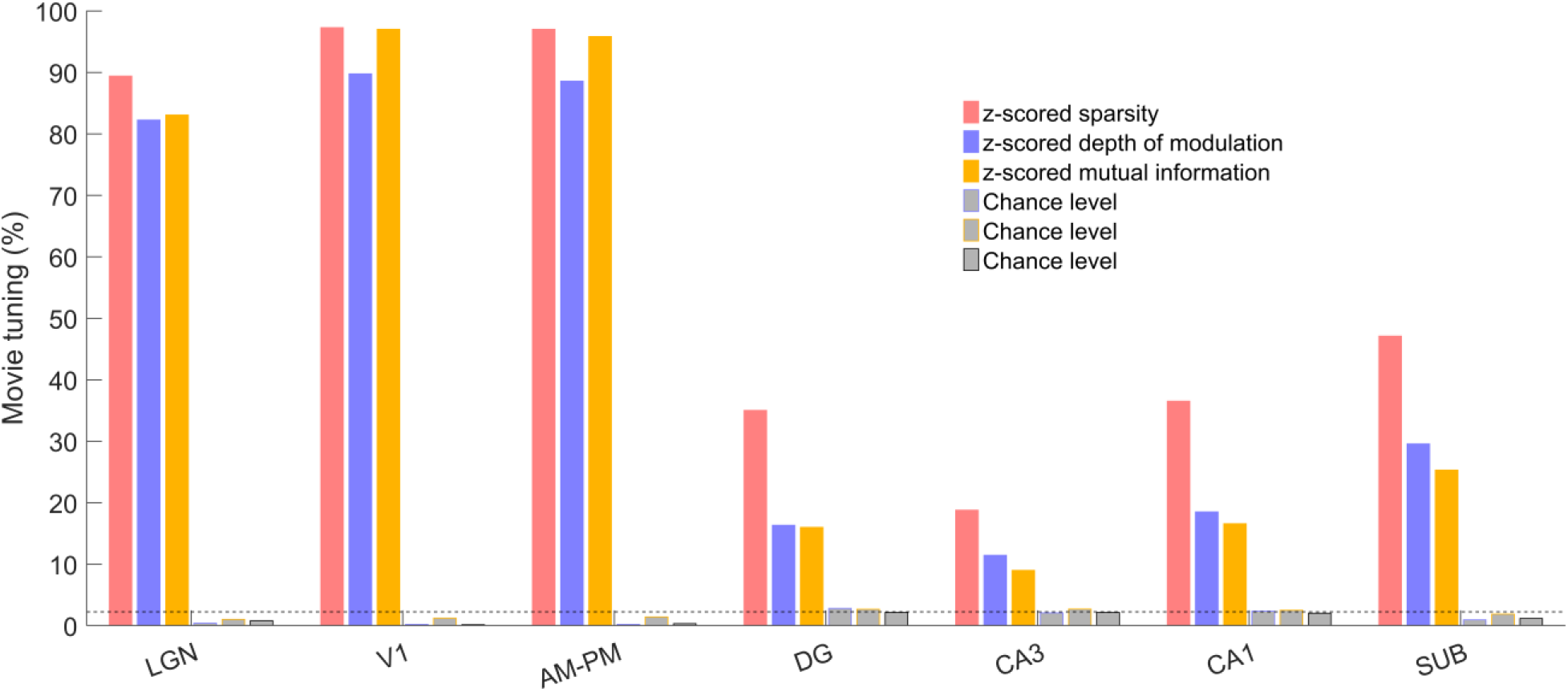
| Multiple metrics show significant and comparable movie tuning. The percentage of movie tuned cells, deemed as z-scored metric > 2, were significantly greater than chance levels (*p*<4.9×10^-11^), using either sparsity or depth of modulation or mutual information as the metric (see *Methods* for metric definitions). Sparsity yielded higher movie tuning than depth of modulation across all brain regions (*p*<1.8×10^-3^), putatively because it captures multi-peaked tuning better than depth of modulation, which only relies on the largest and smallest firing rate responses. Similarly, z-scored mutual information led to greater tuning than chance levels (*p*<4.9×10^-11^), but lesser than that with the sparsity metric (*p*<1.3×10^-5^).

**Figure 1-figure supplement 4.**
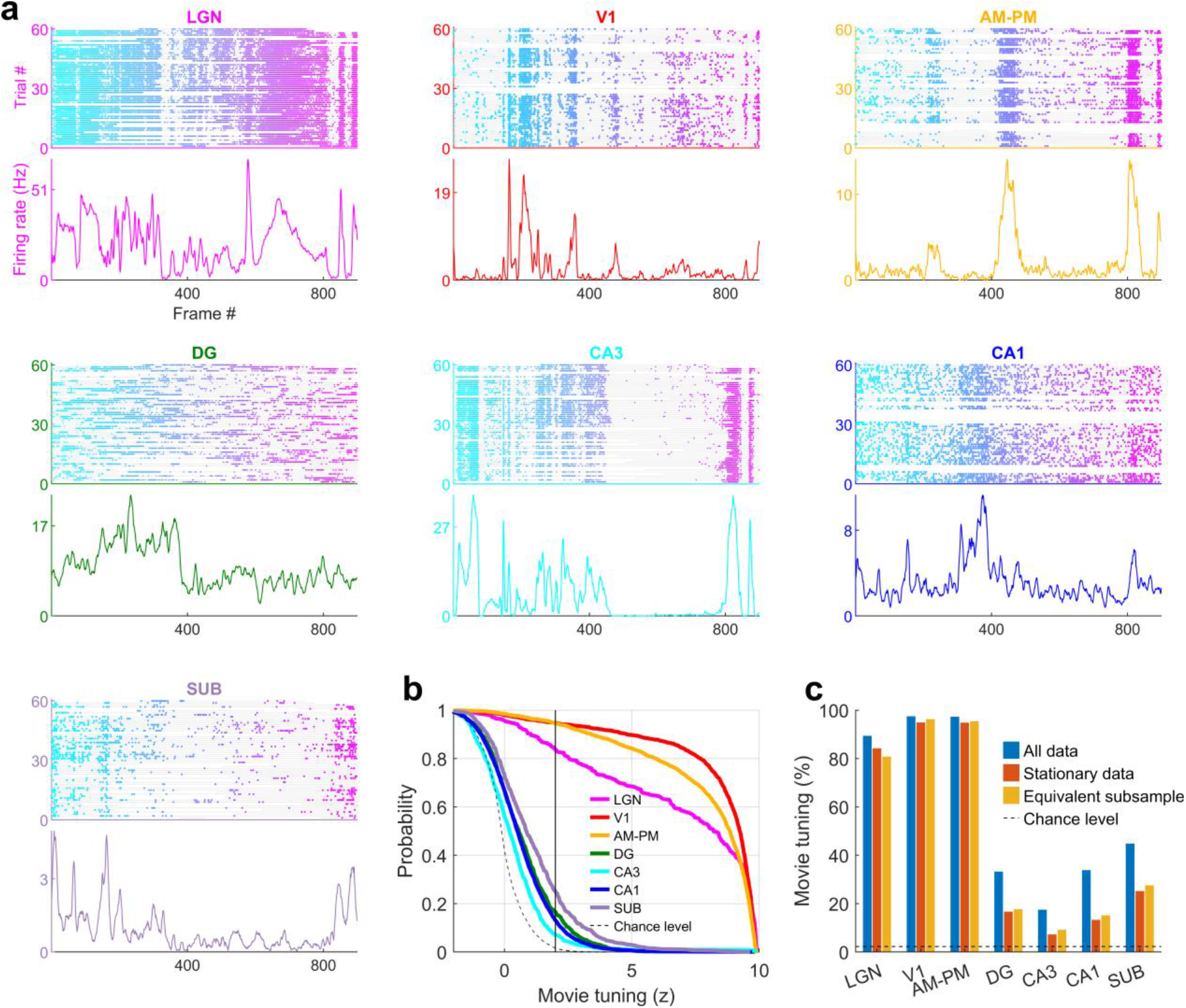
| Movie tuning is intact during immobility. (a) Similar to Figure 1, a representative cell from each of the seven brain regions showing significant modulation movie tuning using only the data when the mouse was immobile, while excluding the data when the mouse was running (stationary data, see *Methods*). **(b)** Fraction of selective neurons was significantly above chance in all brain regions, ranging from 94.7% in V1 up to 7.1% in CA3 in the stationary data. **(c)** To explicitly test the effect of running on movie selectivity, we compared the results in (b) with a random subsample of data, of equal duration as the stationary data, that included running and stationary, to control for the loss of data (see *Methods*). Prevalence of movie selectivity was not significantly different (KS-test *p*>0.05) in these 2 subsamples, except in CA1 (*p*=0.03, 13.1% in stationary data, 15.0% in the equivalent subsample). Only sessions with at least 300 seconds of stationary data were used in this analysis to ensure sufficient statistical power. The reduction in fraction tuned neurons in (b) and (c) for ‘stationary data’, compared to ‘all data’ here and in Figure 1 and Figure 1-figure supplement 2&3 is because of the reduction in the amount of data, which directly reduces statistical significance.

**Figure 1-figure supplement 5.**
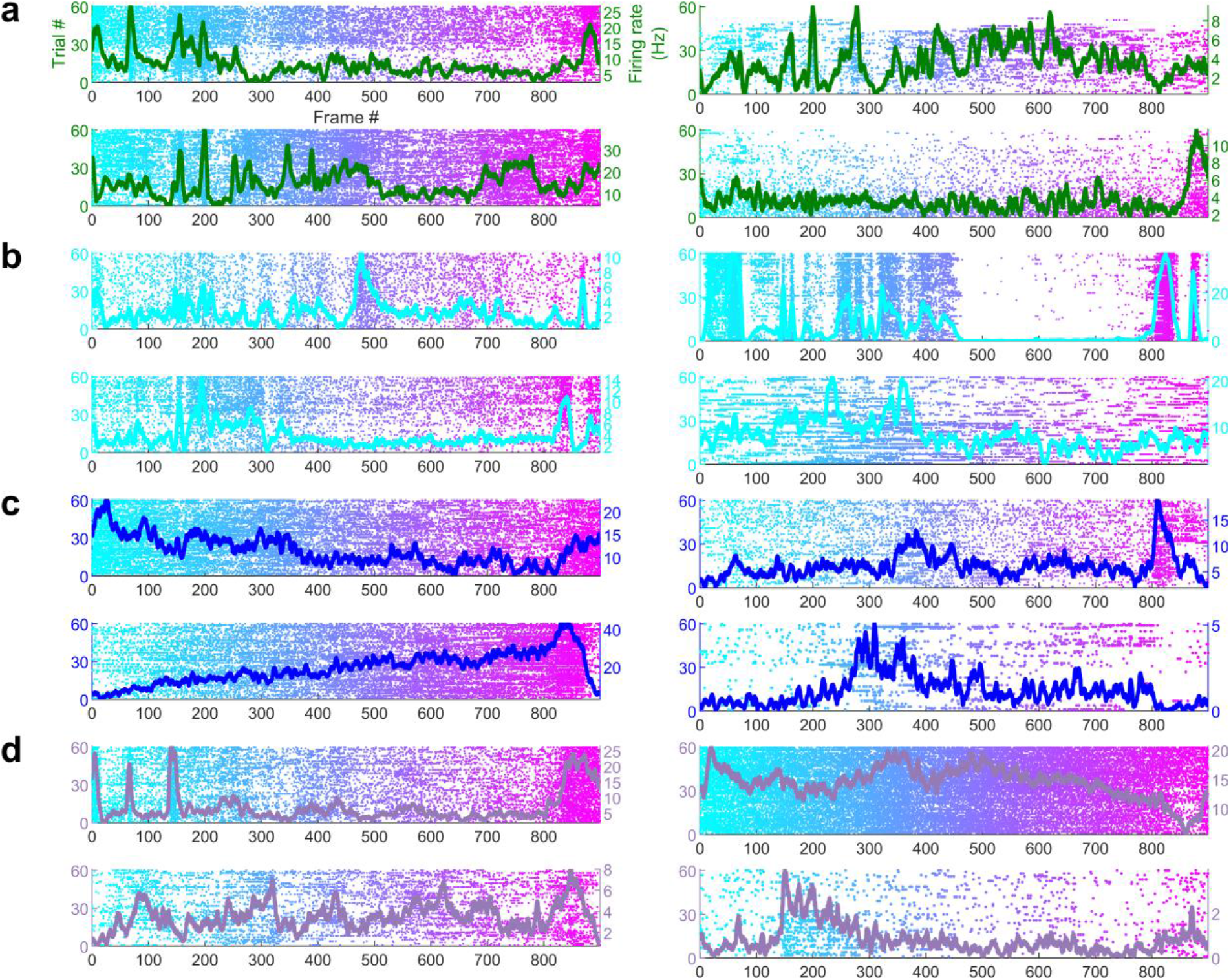
| Simultaneously recorded hippocampal cells have different movie tuning. Four simultaneously recorded and significantly movie-tuned cells each from **(a)** Dentate gyrus, **(b)** CA3, **(c)** CA1 and **(d)** Subiculum. Each cell shows different movie selectivity. Average responses are overlaid (on raster plots), and their color corresponds to the different brain regions, described in Figure 1 legend. This further demonstrates that hippocampal movie tuning is not an artefact of nonspecific variables that alter excitability.

**Figure 1-figure supplement 6.**
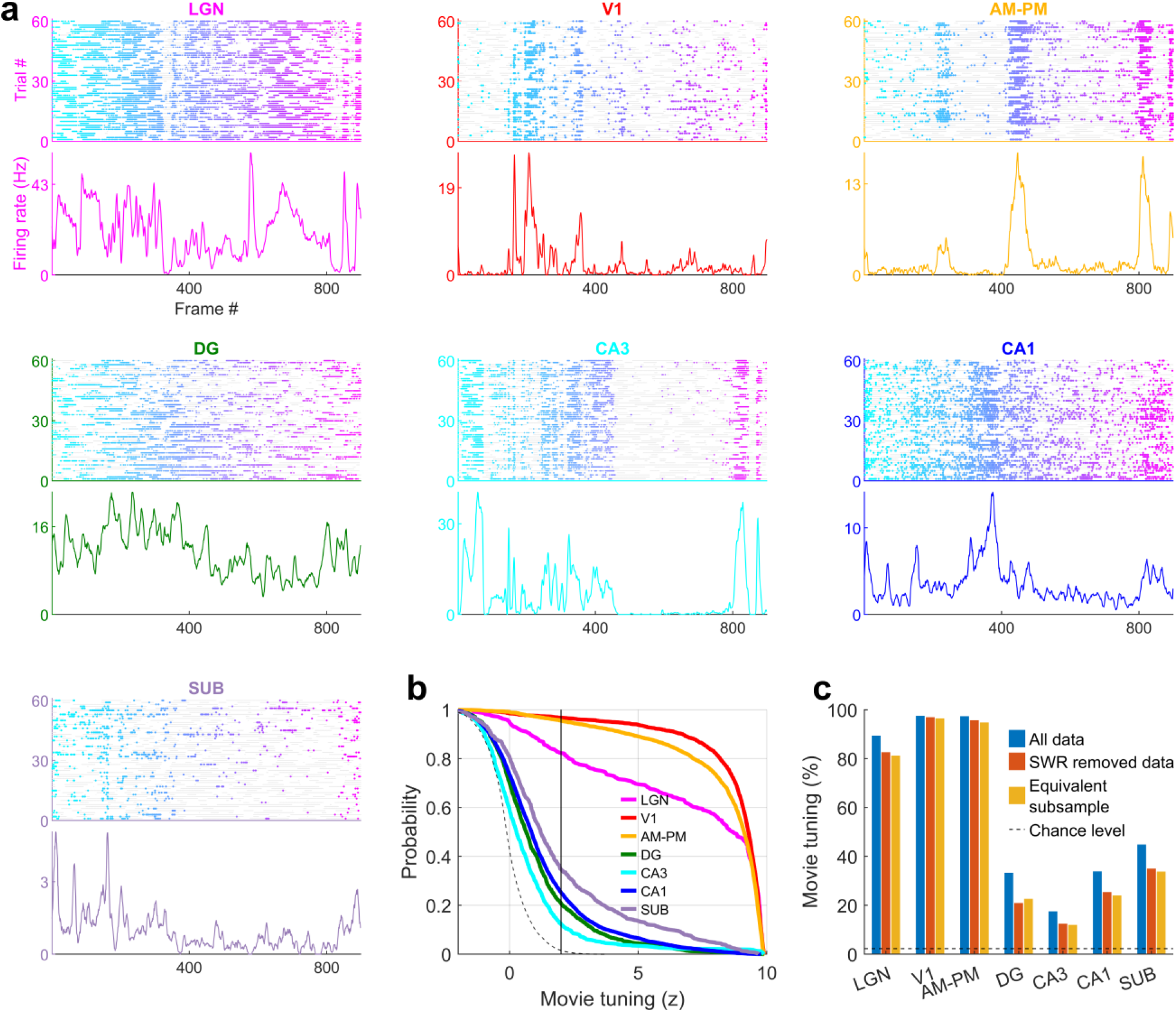
| Movie tuning in unaffected by the removal of sharp-wave ripple (SWR) events. (a) Similar to Figure 1-figure supplement 4, a representative cell from each brain region showing significant modulation movie tuning using data after removal of SWR events (14371 cells from 20 sessions where SWR information was available, see *Methods*). **(b)** Fraction of selective neurons was significantly above chance in all brain regions, ranging from 96.7% in V1 up to 12.3% in CA3 in the SWR removed data. **(c)** To control the loss of data by the removal of SWR, we compared the results with movie tuning in an equivalent subsample of data. Prevalence of movie selectivity was not significantly different (KS-test *p*>0.05) in these 2 subsamples, except in AM-PM (*p*=0.02, 97.1% in SWR removed data, 94.6% in the equivalent subsample). As before (Figure 1-figure supplement 4), due to a reduction in the amount of data, a smaller number of neurons showed significant movie tuning in both SWR removed data as well as equivalent subsampled data.

**Figure 1-figure supplement 7.**
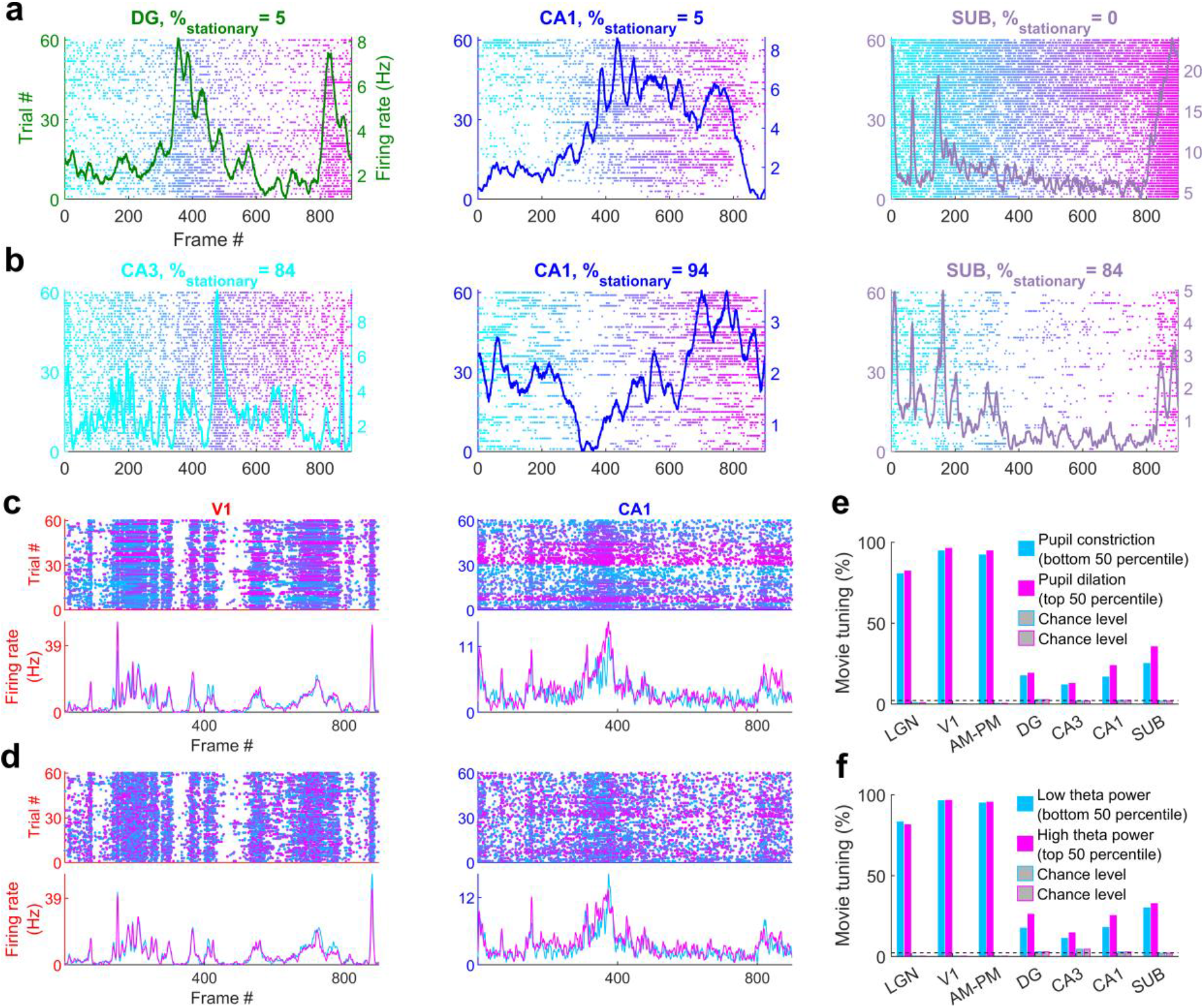
| Movie tuning is comparable across sessions with or without prolonged stationary behavior, high or low pupil dilation or theta power. (a) Three representative cells from sessions with rare stationary periods (% of time in stationary, indicated at the top) and mostly running behavior that showed significant movie tuning**. (b)** Similar to (a), three example cells from sessions with mostly running behavior showing significant movie tuning. Movie tuning persisted across all sessions. **(c)** One representative cell each from V1 and CA1, showing comparable movie tuning during dilated (magenta, high pupil area) or constricted (cyan, low pupil area) pupil. Each dot in the scatter (top) corresponds to one spike, and the color corresponds to the pupil area during that spike. Average movie responses for bottom 50 (cyan-pupil constriction) and top 50 (magenta-pupil dilation) percentiles are shown below. This separation based on 50 percentile ensures equal amount of data in both sub-segments. **(d)** Similar to (c), showing similar movie tuning for data with high (magenta) and low (cyan) theta power. **(e)** Movie tuning in the top as well as bottom 50 percentile of pupil area data was significantly greater than their respective chance levels (*p*<1.2×10^-8^). Top as well as bottom 50 percentile data did not have significantly different movie tuning prevalence for LGN, DG and CA3 (*p*>0.73, which could be because of smaller number of cells recorded in these brain regions), but dilated pupil corresponds to slightly greater tuning for other brain regions (*p*<3.4×10^-4^). **(f)** Similar to (e), the movie tuning in high as well as low theta power data was significantly greater than chance levels (*p*<5.0×10^-10^). Movie tuning was greater in data with high theta power for DG and CA1 (*p*<2.1×10^-6^), but not significantly different for other brain regions (*p*>0.07). Both sub-segments had equal amounts of data to ensure fair comparison.

**Figure 2-figure supplement 1.**
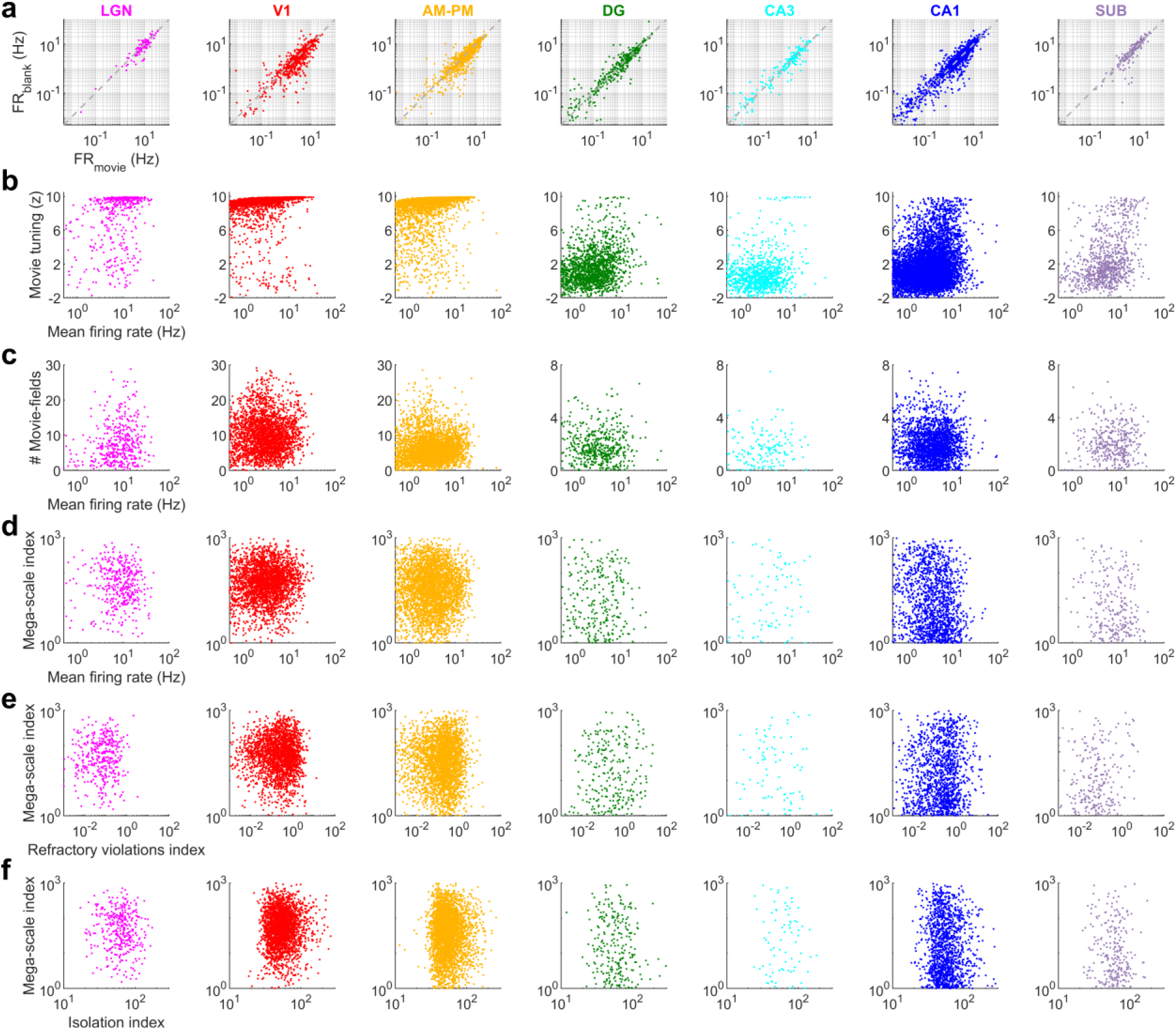
| Few hippocampal neurons had greater than 5 movie-fields. Only a handful of movie-tuned neurons from dentate gyrus (row 1 and 2), CA3 (row 3 and 4), CA1 and subiculum (bottom-right), had more than 5 distinct movie-fields. Similar format as Figure 1 and Figure 1-figure supplement 5. This is in contrast to the visual areas where a large number of movie fields were common and the average number of movie fields per cell was greater than 6 (Figure 2).

**Figure 2-figure supplement 2.**
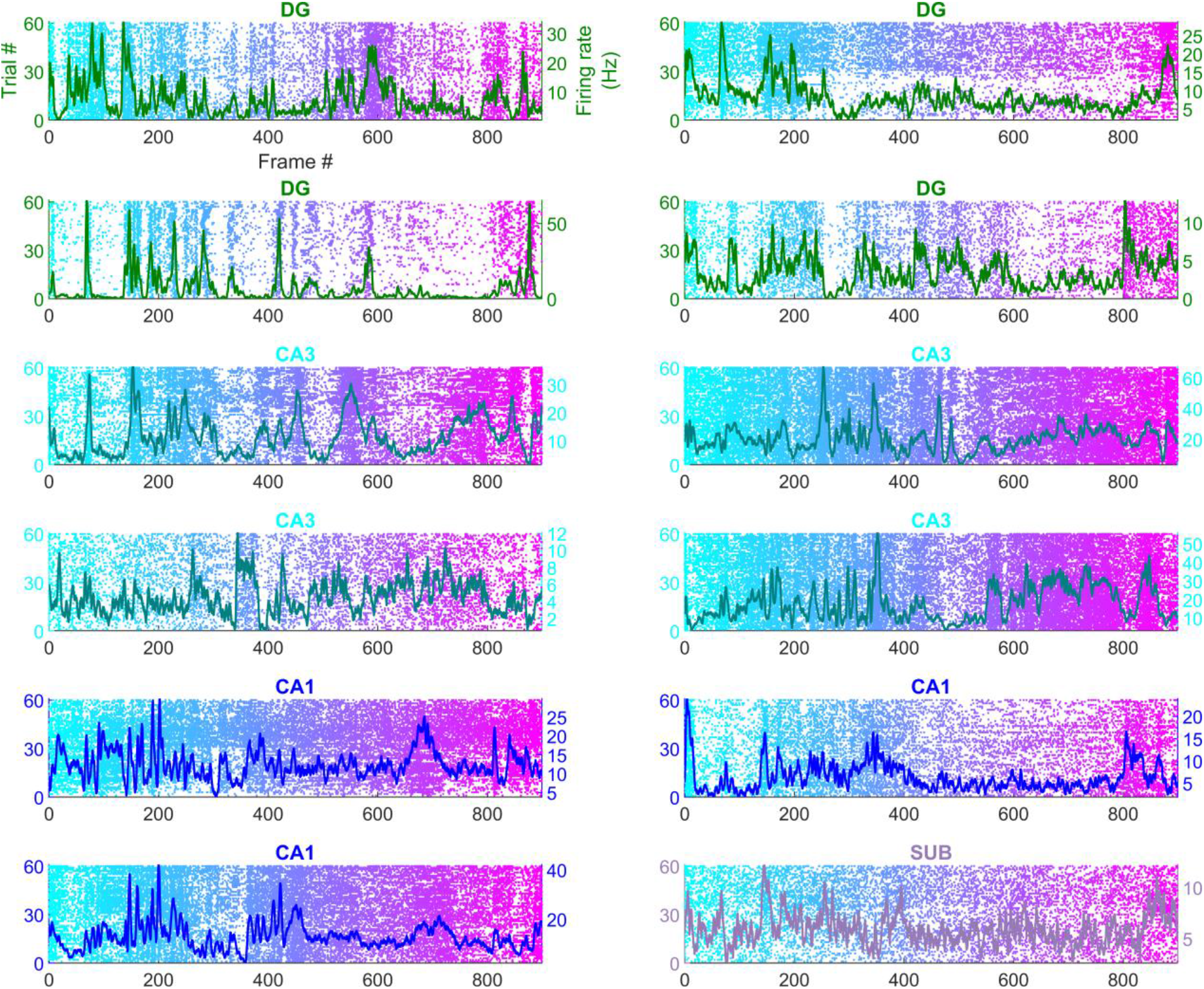
| Mega-scale movie-coding within a single cell is smaller than the ensemble wide mega-scale index in visual, but not hippocampal areas. (a) Distribution of median duration of movie-fields, computed across all fields of a single neuron. Median movie-field duration was significantly larger in all hippocampal areas compared to all visual areas (KS-test *p*<7.1×10^-31^). Median field duration between DG-CA3 and DG-CA1 were not significantly different, but all other visual-visual and hippocampal-hippocampal region pairs were significantly different. (KS-test *p*<0.04). CA3 had the largest median field duration (6.3±0.48s), and V1 had the smallest (0.27±0.03sec). Surprisingly, LGN movie-field durations (0.57±0.13s) were about two-fold longer than V1 (*p*=2.5×10^-21^); though both were smaller than those in the higher order brain areas (0.71±0.05s). **(b)** Firing in movie-fields, normalized by that in the shuffled response were used to obtain the median value from all fields of a neuron. This metric of median movie-field activation is significantly different across all brain region pairs (KS-test *p*<3.4×10^-5^), except DG-CA3, CA3-CA1 and DG-CA1 pairs. The largest median movie-field activation was in V1 (2.5±0.05), and the smallest in subiculum (1.13±0.03). **(c)** Cumulative firing in movie-fields, normalized by that in the shuffle response, obtained by adding the activity within all fields of a neuron was significantly different across all brain region pairs (KS-test *p*<3.0×10^-7^), except DG-CA3, CA3-CA1 and DG-CA1. V1 response was largest (1.93±0.04), and subiculum was the smallest (1.11±0.02). **(d)** For each brain region, the movie-field duration ratio was recalculated by randomly reassigning the cell ids to all the movie peaks from that brain region. Using this new assignment of movie peaks to a cell, we obtain the expected mega-scale index (largest/smallest peak duration) based on the ensemble behavior. The observed mega-scale index within a cell was smaller than expected from the ensemble in all the visual areas (KS-test *p*<3.2×10^-3^, median was 77.5%, 56.2% and 41.7% of chance for LGN, V1 and AM-PM respectively). This was not the case in hippocampal regions (*p*>0.23). Thus, individual cells in the visual, but not hippocampal, areas sampled a subset of possible mega-scale coding values of the ensemble. **(e)** Histogram of movie-fields, binned for their durations (log-scaled) and their prominence (also log-scaled). The most prominent fields tended to be wider in most brain areas, and this effect was stronger in hippocampal regions, than visual. Note that the histogram color is also log-scaled.

**Figure 3-figure supplement 1.**
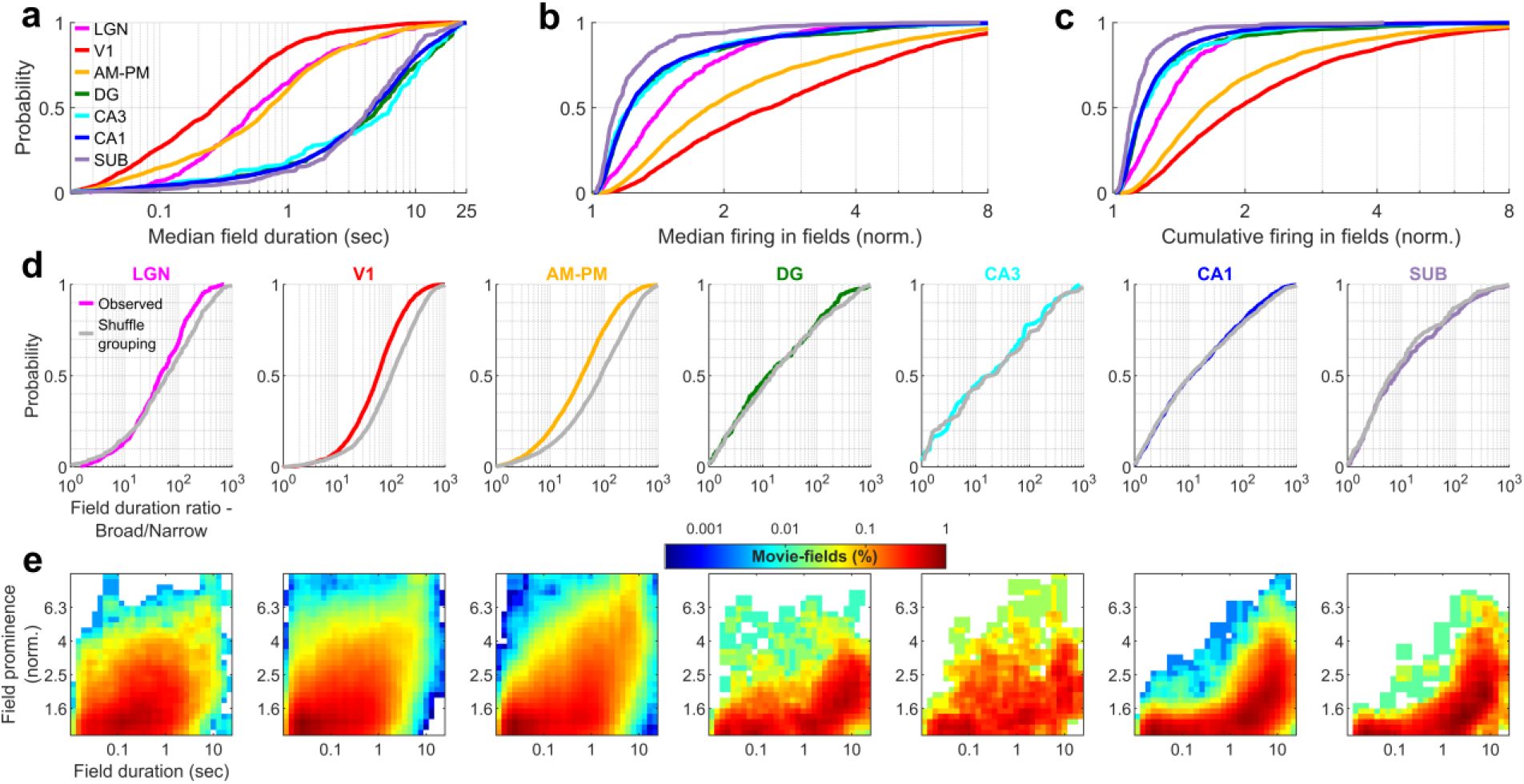
| Population vector overlap is wider in hippocampus than visual areas. **(a)** Population vector overlap between even and odd trials for the population of tuned neurons show highest overlap along the diagonal (i.e. for the same movie frame) for all brain regions. Each neuron’s response was normalized by its mean rate and the average response in even as well as odd trials was smoothed by a Gaussian window of 2 frames (66.6ms, see *Methods*). Dashed black lines indicate the - 300 and +300 frames away from the diagonal. Notice large correlations (close to unity, horizontal color bar) indicating stable responses. The correlations decay quickly to smaller values for the visual areas but more slowly for hippocampal areas, due to their broader movie-fields. **(b)** Same as (a), but for untuned neurons, resulting in a salt and pepper overlap pattern and low values of correlation, indicating lesser stability than the tuned neurons. Since the majority of cells in the visual areas were tuned, the untuned population was smaller, leading to more variable population vector overlap. **(c)** The average population vector overlap, computed across all frames, as a function of the number of movie frames away from the diagonal in (a). It had a large value in visual regions for the 0^th^ diagonal (colored lines) indicating stable responses, whereas the untuned neuron population (gray lines) were unstable, with values near zero, or chance level. The highest population vector overlap in hippocampal regions was smaller than visual areas but persisted for more frames, due to their broader movie-fields (Full width at half maximum of the peak – 17.3 frames for LGN, 22.7-V1, 39.0-AM&PM, 49.8-DG, 57.4-CA3, 64.7-CA1 and 59.2-subiculum).

**Figure 3-figure supplement 2.**
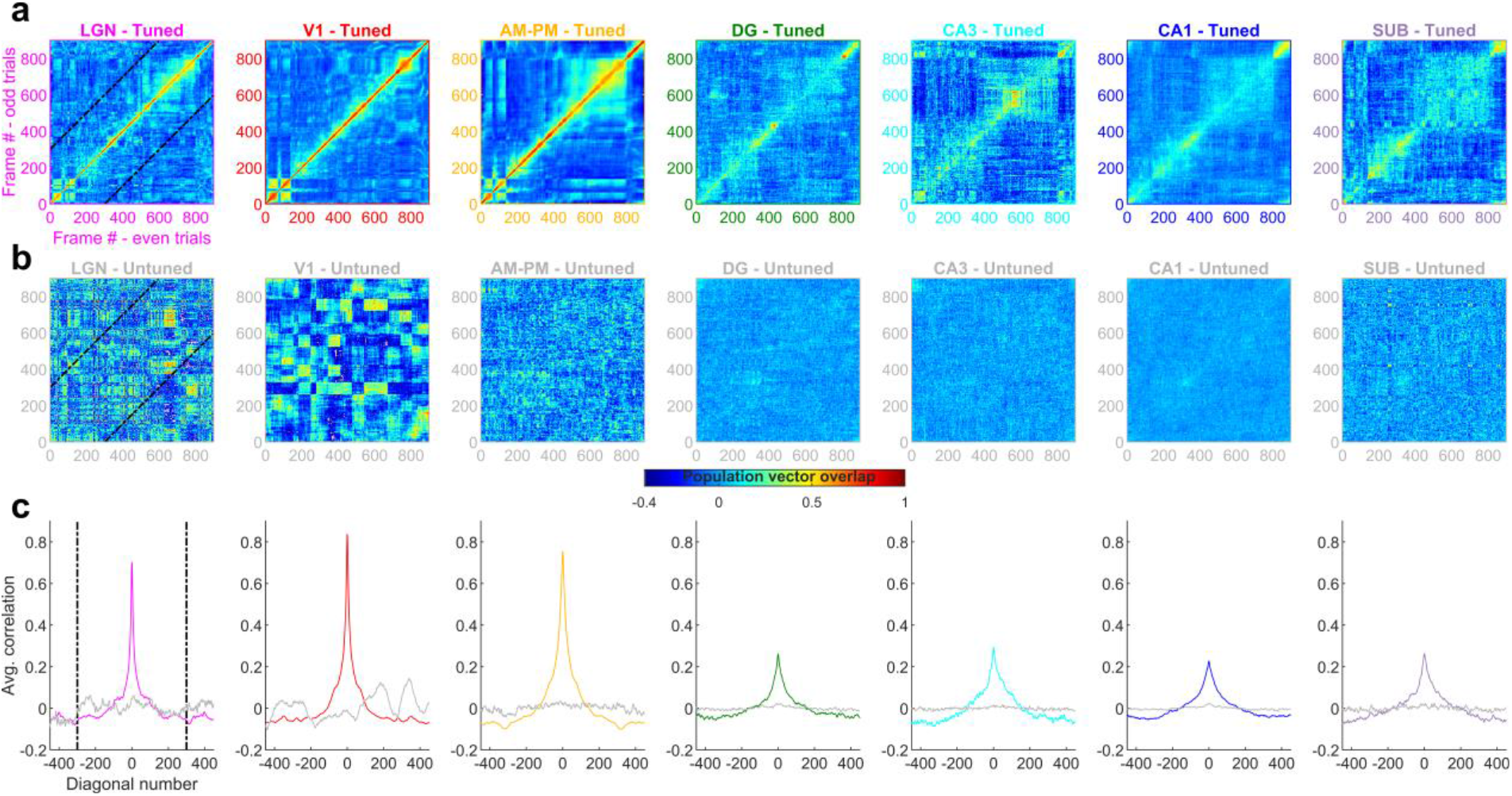
| Movie-field properties strongly reflect the frame-to-frame correlation structure of the movie in the visual but not hippocampal areas. (a) The adjacent movie frame (frame_n_,frame_n+1_) correlation coefficient, indicating the similarity of 2 consecutive frames, termed F2F image correlation, is shown in gray. Similarly, the correlation coefficient between the population vector of neural responses between adjacent frames, was termed F2F neural correlation, computed separately for each brain region is shown in color. The relationship between F2F-image, and F2F-neural correlation across brain regions is shown in the matrix on the right. Diagonal entries indicate correlation between F2F-image and F2F-neural, with largest correlation for LGN (+0.82), followed by V1 (+0.75), CA3 (+0.56), DG (+0.52), AM&PM (+0.38), CA1 (+0.14) and SUB (+0.07). All correlations were significant (*p*<1.1×10^-3^). Above diagonal entries indicate the correlation coefficient between brain region pairs. Below diagonal entries indicate the same but using partial correlations that factor out the F2F image correlation. All partial correlations were significant (*p*<5.4×10^-3^), except V1-CA1 and AM&PM-CA3. **(b)** Histogram of the number of movie-field peaks (i.e. movie field density) across all tuned neurons in a brain region, as a function of the movie frame. This distribution was significantly non-uniform (Chi-square goodness-of-fit test for uniform distribution, *p*<3.8×10^-6^) for all brain regions. All distributions were significantly negatively correlated with F2F image correlation (*p*<10^-7^). These correlations were much stronger in visual (LGN -0.77, V1 -0.73, AM-PM -0.71) areas than hippocampal areas (DG -0.23, CA3 -0.70, CA1 -0.18, SUB -0.27). The largest partial correlation after factoring out the F2F image correlation was between LGN-V1 and V1-CA3 (0.89) and the least between LGN-DG (0.16). All partial correlations were significant (*p*<1.2×10^-6^). **(c)** Same as (b), but for the median duration of movie-fields. F2F image correlation shown in gray, with larger correlation between consecutive frames between frames 400-800 clearly reflected in larger movie-field durations in visual areas. All distributions were significantly non-uniform (Chi-square goodness-of-fit test, *p*<10^-100^) and all distributions were significantly positively correlated with F2F image correlation (*r*>0.24, *p*<2×10^-13^), with greater values for visual areas (LGN +0.61, V1 +0.51, AM-PM +0.55) than hippocampal (DG +0.39, CA3 +0.58, CA1 +0.42, SUB +0.24). Note that the y-axes for the histogram are log-scaled and show larger median durations for hippocampal regions than visual. The largest correlation was between AM&PM-CA3 (0.81) and the least between CA3-SUB (−0.03). All partial correlations were significant (*p*<4.4×10^-4^), except LGN-CA1 (*p*=0.11) and CA3-SUB (*p*=0.39). **(d)** Total firing rate across all broad spiking neurons in different brain regions, showing similar non-uniformity as Figure 3c. All brain regions had significantly negative correlation with the F2F image correlation (*r*<-0.08, *p*<0.03), except DG, which was significantly positively correlated (*r*=0.21, *p*=2.4×10^-10^). The largest number of above chance (gray lines) deviations were seen for AM-PM (340 frames), and least for CA3 (57 frames), which could be due to the low cell count in CA3. Below chance level deviations were least common in LGN (25 frames), and most common in AM-PM (441 frames). The largest partial correlation amongst brain region pairs was between DG-CA1 (0.76) and the least between V1-DG (−0.1) after factoring out the F2F image correlation. All partial correlations were significant (*p*<3.2×10^-3^), except LGN-DG (*p*=0.81). Similar to Figure 3c-e.

**Figure 4-figure supplement 1.**
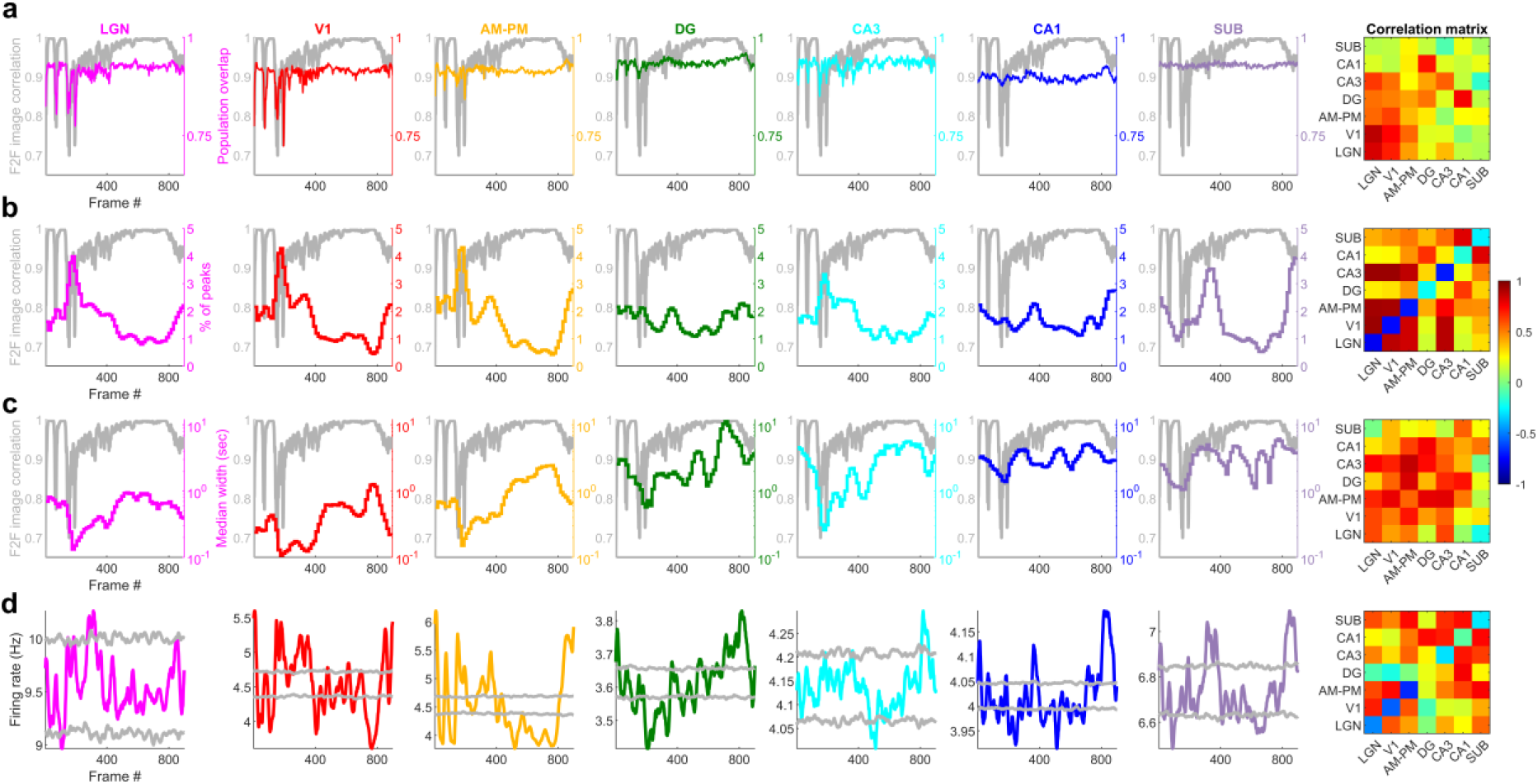
| Scrambled movie elicits narrower but more movie-fields per cell than the continuous movie in all the visual regions. (a) Cumulative distribution of the total number of fields per cell for the scrambled movie shows the largest number of fields in LGN (mean±s.e.m., 31.8±2.0), followed by V1 (24.0±0.38) and last AM-PM (11.1±2.1). All three brain regions were significantly different from each other (KS-test *p*<2.0×10^-5^). **(b)** The median scrambled movie-field duration was shortest in LGN (43.9±131.2ms), intermediate in V1 (46.2±24.8) and widest in AM-PM (77.6±40.1ms), and differences were significant (*p*<7.0×10^-4^). This was much smaller than for the continuous movie (Figure 2). **(c)** Durations of fields for scrambled sequence across all fields of all neurons from a brain region. These were narrowest in LGN (31.3±6.5ms), followed by V1 (38.6±0.2) and last AM-PM (64.3±6.7). All differences were significant (KS-test *p*<7.2×10^-136^). **(d)** Despite these differences, the cumulative duration of movie-fields was comparable across the three brain regions (1.69±0.05sec for V1, 2.03±0.07 for AM-PM and 2.4±0.2 for LGN), but significantly different (*p*<1.7×10^-5^). Note the linear scale on the x-axis in this panel compared to the log-scale in other panels. **(e)** Ratio of field durations, i.e., mega-scale index, for the scrambled movie was smallest in V1 (15.5±1.6), intermediate in LGN (16.3±5.4) and largest in AM-PM (23.4±2.1), and not significantly different between V1 and LGN (*p*=0.28). V1-AM&PM and LGN-AM&PM were significantly different (*p*<5.7×10^-5^). **(f)** Cumulative spiking activity, summed across all movie-fields of a given neuron was largest in V1 (3.8±0.1), intermediate in LGN (2.3±0.1) and smallest in AM-PM (2.0±0.07), and significantly different between all brain region pairs (*p*<0.02).

**Figure 4-figure supplement 2.**
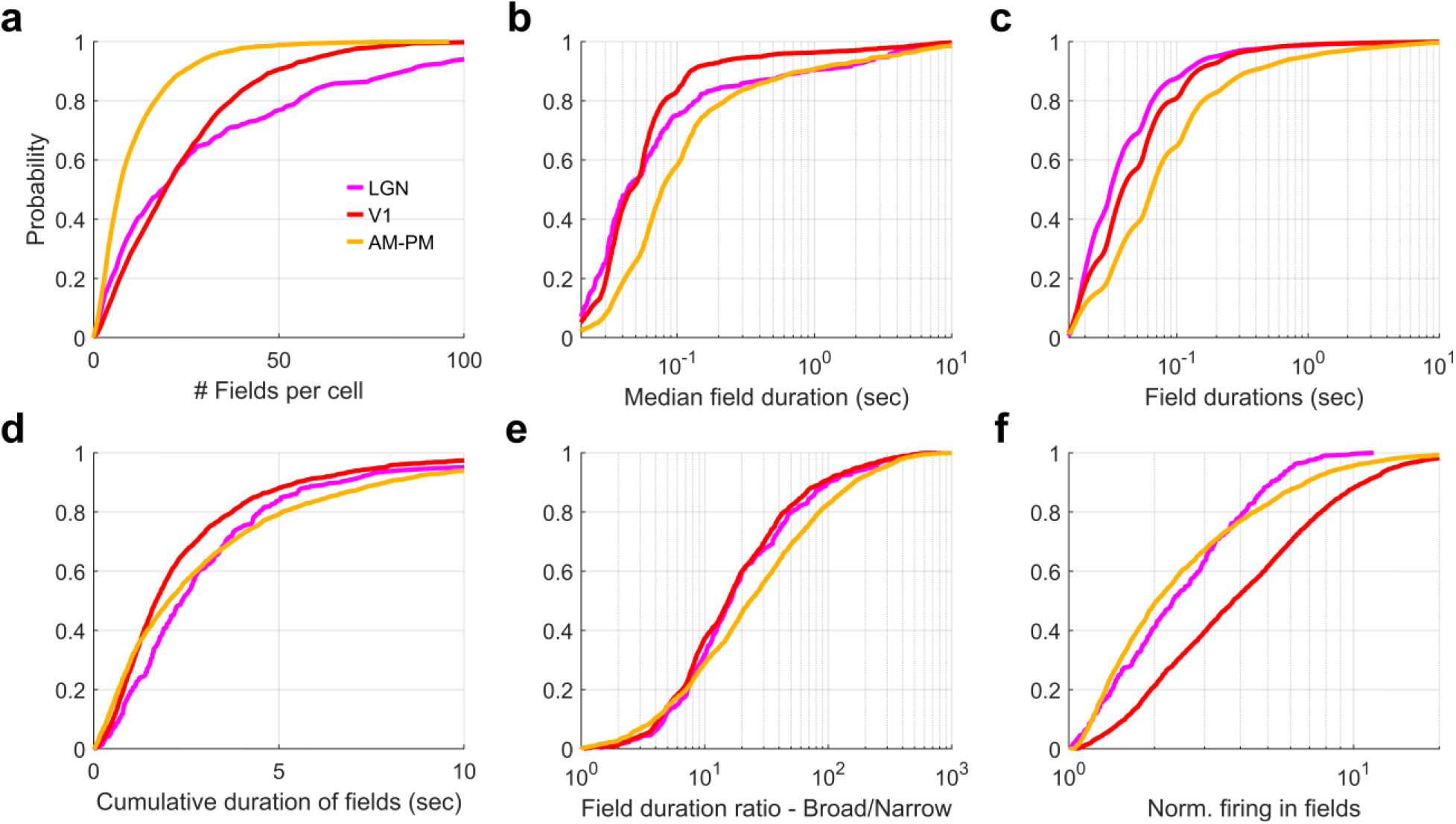
| Cell by cell comparison of continuous vs scrambled movie responses. Data for only those visual area neurons that were significantly modulated by both the continuous and scrambled movie were used. **(a)** The number of movie-fields per cell for the continuous movie was significantly smaller than that for scrambled sequence in all brain areas (LGN – continuous mean±s.e.m.=10.7±0.42, scrambled=31.8±2.0, KS-test *p*=2.0×10^-23^, V1-10.8±0.11 vs. 24.0±0.38, KS-test *p=*3.7×10^-210^, AM&PM-6.9±0.07, vs. 11.1±0.21, KS-test *p*=1.3×10^-57^). Data are additionally scattered by a small random number for the ease of visualization. **(b)** Median duration of movie-fields for a cell was significantly larger for continuous movie, compared to scrambled sequence in all visual regions. (LGN continuous=0.46±0.08s, scrambled=0.04±0.13s, KS-test *p*=7.1×10^-65^, V1-0.25±0.03s vs. 0.04±0.02s, KS-test *p<*10^-150^, AM&PM 0.65±0.04s, vs. 0.08±0.04s, KS-test *p*<10^-150^). **(c)** Cumulative duration of all movie-fields for a cell was significantly larger for continuous movie, compared to scrambled sequence in all visual regions. (LGN continuous=8.9±0.23s, scrambled=2.4±0.19s, KS-test *p*=3.2×10^-69^, V1-6.1±0.09s vs. 1.69±0.05s, KS-test *p=*3.3*x*10^-296^, AM-PM 7.8±0.1s, vs. 2.0±0.07, KS-test *p*=9.0×10^-318^). **(d)** Histogram of number of fields per cell, for continuous and scrambled movies. **(e)** Logarithmically spaced histogram of median field durations was significantly different between continuous and scrambled sequence. **(f)** Similar to (e), histogram of cumulative duration of movie-fields for each cell. **(g)** The ratio of number of fields per cell between continuous and scrambled movies was biased to smaller than unity values for all brain regions, with the largest bias for LGN (0.46±0.08), intermediate for V1 (0.5±0.04), and least for AM-PM (0.77±0.05). **(h)** The median field duration ratio was biased to values greater than unity, with the largest bias for LGN (7.4±1.4), least for V1 (4.5±0.68), and intermediate for AM-PM (5.5±0.82). **(i)** The cumulative field duration ratio was also biased to values greater than unity, with similar biases for LGN (3.37±0.36), V1 (3.1±0.3), and AM-PM (3.3±0.67).

**Figure 4-figure supplement 3.**
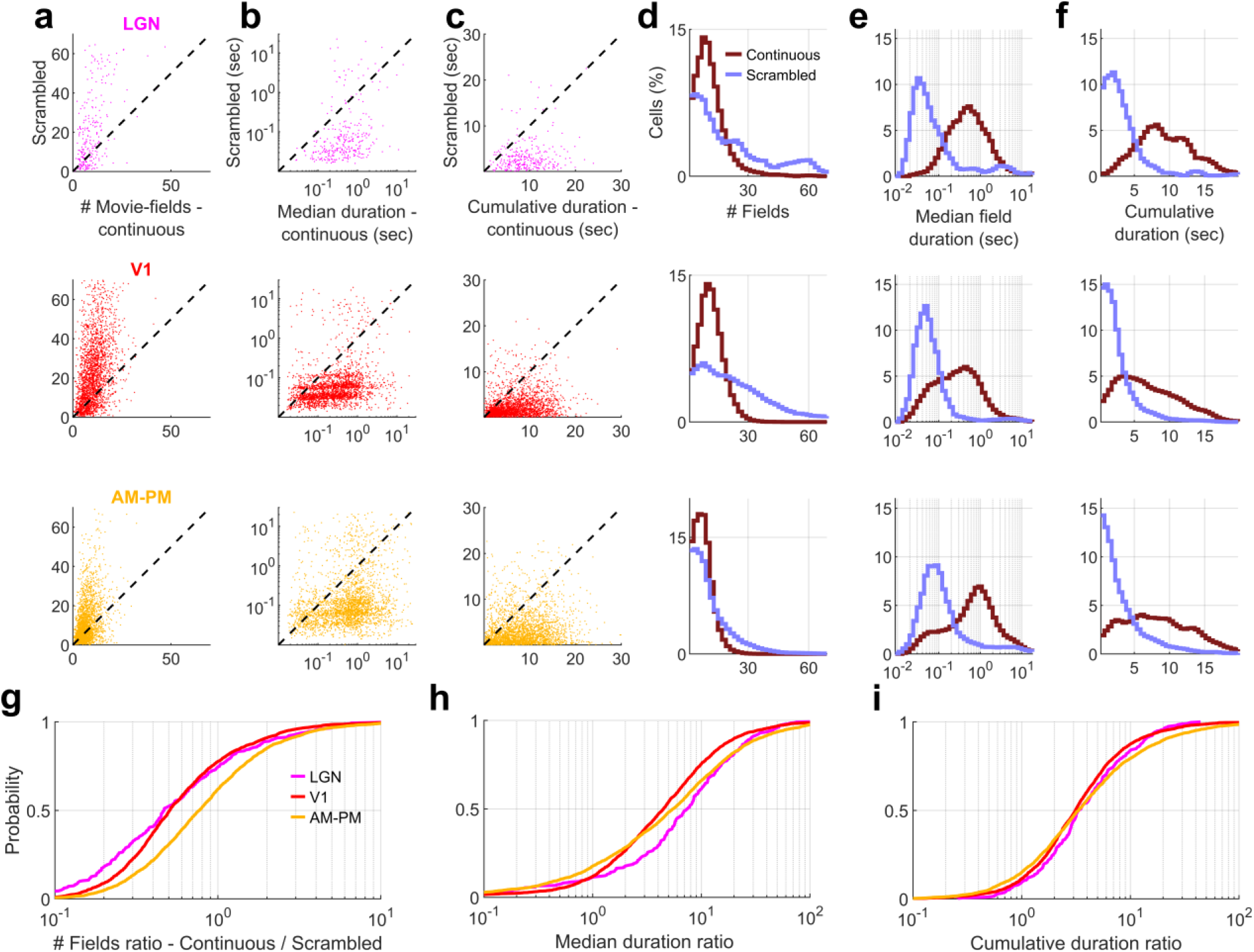
| Multiple-single unit activity (MSUA) across all movie-tuned neurons in a brain region shows greater modulation than chance for the scrambled sequence in all visual areas. (a) Stack plot of tuned responses to the scrambled movie presentation from each brain region, sorted according to the frame with peak response. Each firing rate profile is normalized by the peak response causing the diagonal to be unity. The average firing rate of non-peak frames (similar to Figure 3b, legend) was smallest (0.50*x* of the average peak response across all neurons) for V1, followed by AM&PM-0.56 and largest for LGN-0.65. **(b)** Colored trace-average response, across all tuned responses from (a). gray trace - chance level, z=±4, corresponding to the *p*=0.025 level after Bonferroni correction. **(c)** Number of frames for which the observed response exceeds (or falls below) z=±4 cutoff from (b), called significantly deviant frames. V1 had the largest number of positive (279 frames) and negative (297) deviant frames, similar to the continuous movie (Figure 3d, 289 positive and 324 negative). AM-PM had intermediate (225 & 235) deviant frames for the scrambled movie, which was lower than the continuous movie (285 & 454). LGN had the least number of significantly deviant frames (31 & 29), larger than the continuous movie (12 & 0). **(d)** Firing rate deviation above chance levels, corresponding to the significant frames, as identified in (c), normalized by the mean rate of the MSUA. Largest deviation was observed in V1 (above-3.1 and below-2.7%), and least in LGN (1.1% and 0.45%) Compare with Figure 3. **(e)** Frame to frame (F2F) image correlation, from Figure 4a for comparison. This had no structure and values hovered around zero (unlike the continuous movie) and was not significantly correlated with the MSUA responses in (b), for any of the brain regions (*Pearson* correlation coefficient LGN *p*=0.71, V1 *p*=0.06, AM-PM *p*=0.21). Despite this, the MSUA shows significant modulation.

**Figure 4-figure supplement 4.**
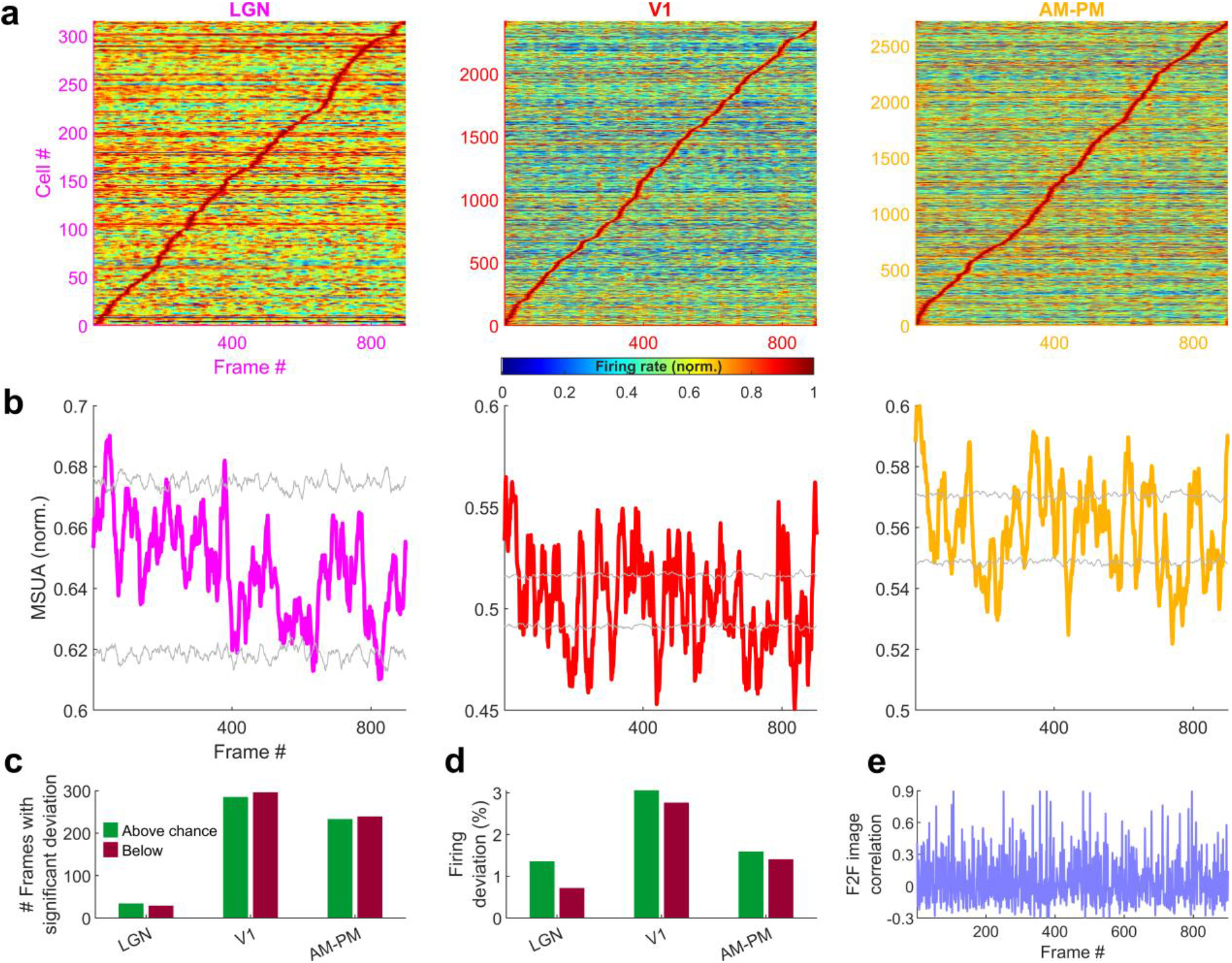
| Latency of responses to the scrambled-sequence corresponds to the anatomical hierarchy of visual areas. (a) Average response for one representative cell from each visual region, that had high similarity between the continuous movie and the rearranged scrambled sequence responses (see *Methods*). Gray response in background corresponds to the chronological scrambled sequence. **(b)** Cumulative histogram of z-scored correlation between continuous and scrambled-rearranged tuning responses (see *Methods*). Dotted black line indicates significance threshold of z>2. **(c)** The latency at which continuous and scrambled-rearranged responses were maximally correlated showed high values (heuristically above 0.25) in a short range of positive latencies for LGN, V1 and AM-PM neurons. This analysis was restricted to neurons tuned in continuous as well as scrambled movies. Similar analysis for hippocampal regions resulted in almost no correlations above 0.25. **(d)** Cumulative histogram of latencies, when the continuous and scrambled-rearranged responses were maximally correlated, was the smallest for LGN (59.5±4.6ms), and largest for higher visual areas, AM-PM (91.6±1.6ms). Hippocampal regions were excluded, owing to lack of data with correlation above 0.25.

**Figure 4-figure supplement 5.**
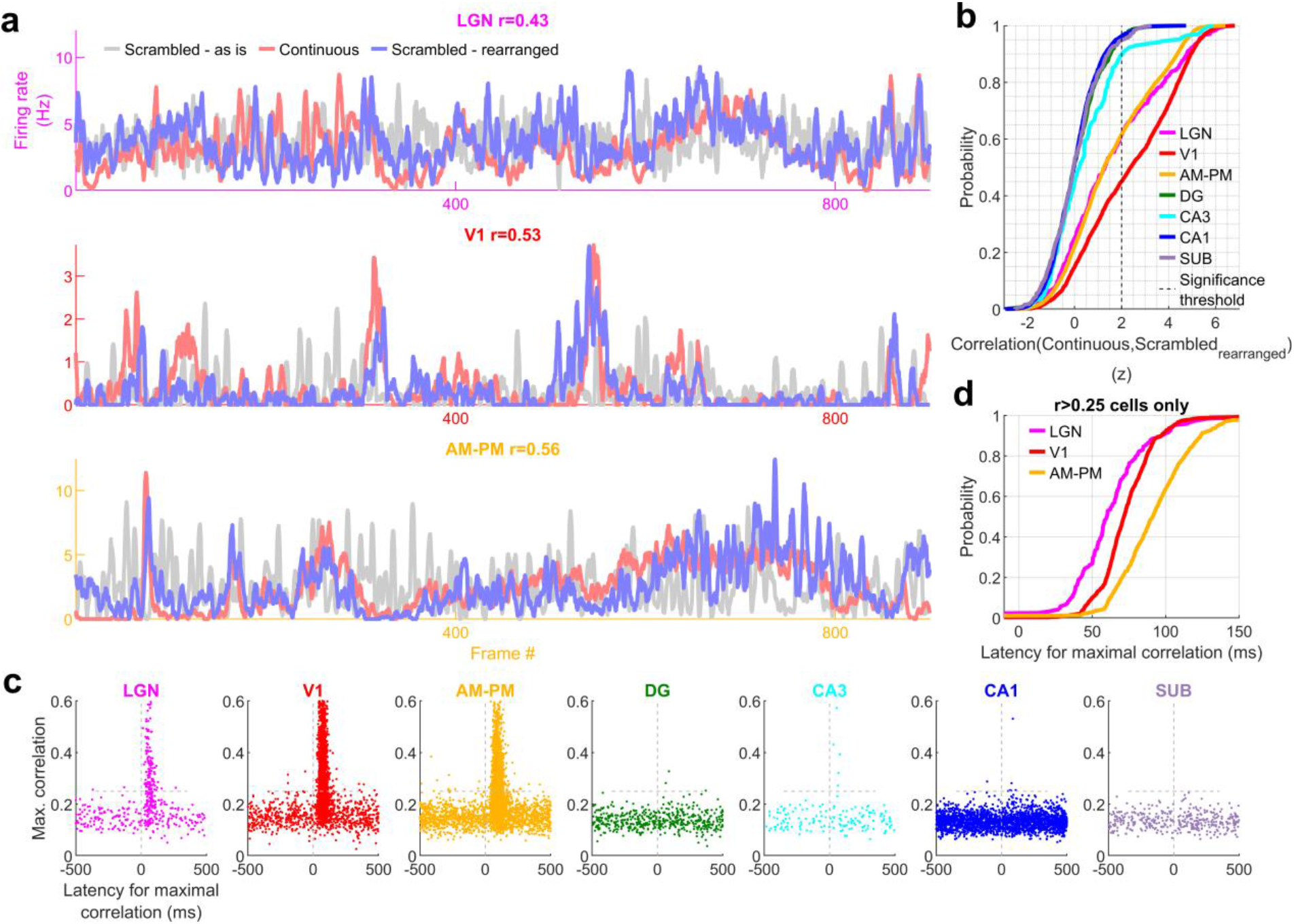
| Movie tuning in hippocampal neurons remains near chance level even after rearranging scrambled movie frames. Histogram showing percentage of tuned cells for movie presentation in the continuous (red), scrambled order, taken as is (light blue), or the scrambled order but rearranged (dark blue). Movie tuning was significantly higher for the continuous presentation (*p*<3.5×10^-3^) than the scrambled as is condition or scrambled rearranged condition (*p*<2.6×10^-6^), in all brain regions. Movie tuning for the scrambled presentation taken as is, or after rearrangement was not significantly different for all brain regions (*p*>0.08), except LGN (*p*=1.3×10^-5^) and V1 (*p*=0.001), although the prevalence of tuning was comparable (63.7 and 64.3%-LGN and 90.1 and 90.0%-V1).

**Figure 4-figure supplement 6.**
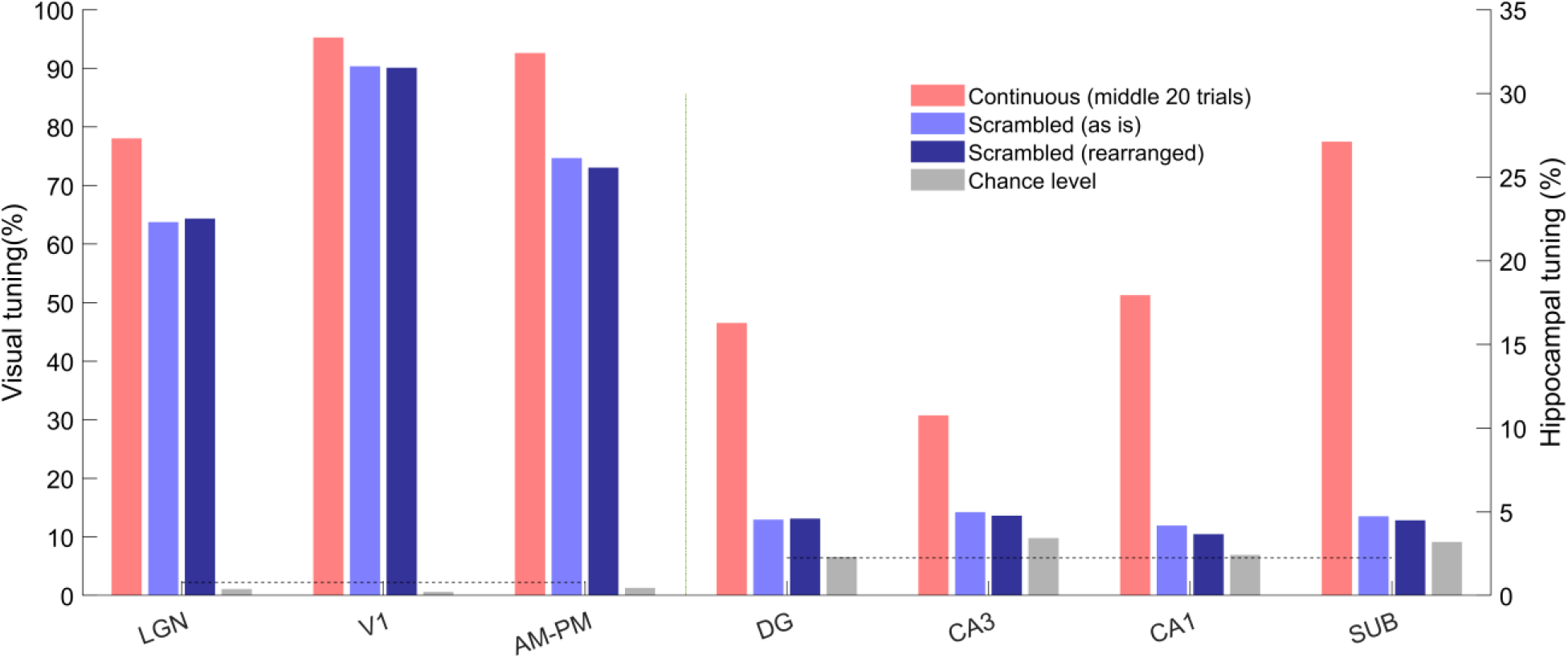
| Population vector overlap was narrower for the scrambled compared to the continuous movie. (a) Population vector overlap between even and odd trials for tuned neurons showing higher overlap along the diagonal for all brain regions. Black lines indicate the -300 and +300 diagonal, whereas the main diagonal is the 0^th^ diagonal. **(b)** Same as (a) but for untuned neurons, resulting in a salt and pepper overlap without higher correlation around the diagonal. **(c)** The average overlap along diagonals had a large value in visual regions for the 0^th^ diagonal, which was not true for the untuned neuron population. Average correlation in hippocampal regions was broader and lesser in magnitude compared to visual regions. Similar to Figure 3-figure supplement 1. Full width at half maximum of the peak – 4.4 frames for LGN, 4.8-V1, 5.2-AM&PM, 7.6-DG, 5.7-CA3, 10.8-CA1 and 15.1-subiculum, even though consecutive frames in the scrambled presentation were largely uncorrelated.

**Figure 1-figure supplement 8.**
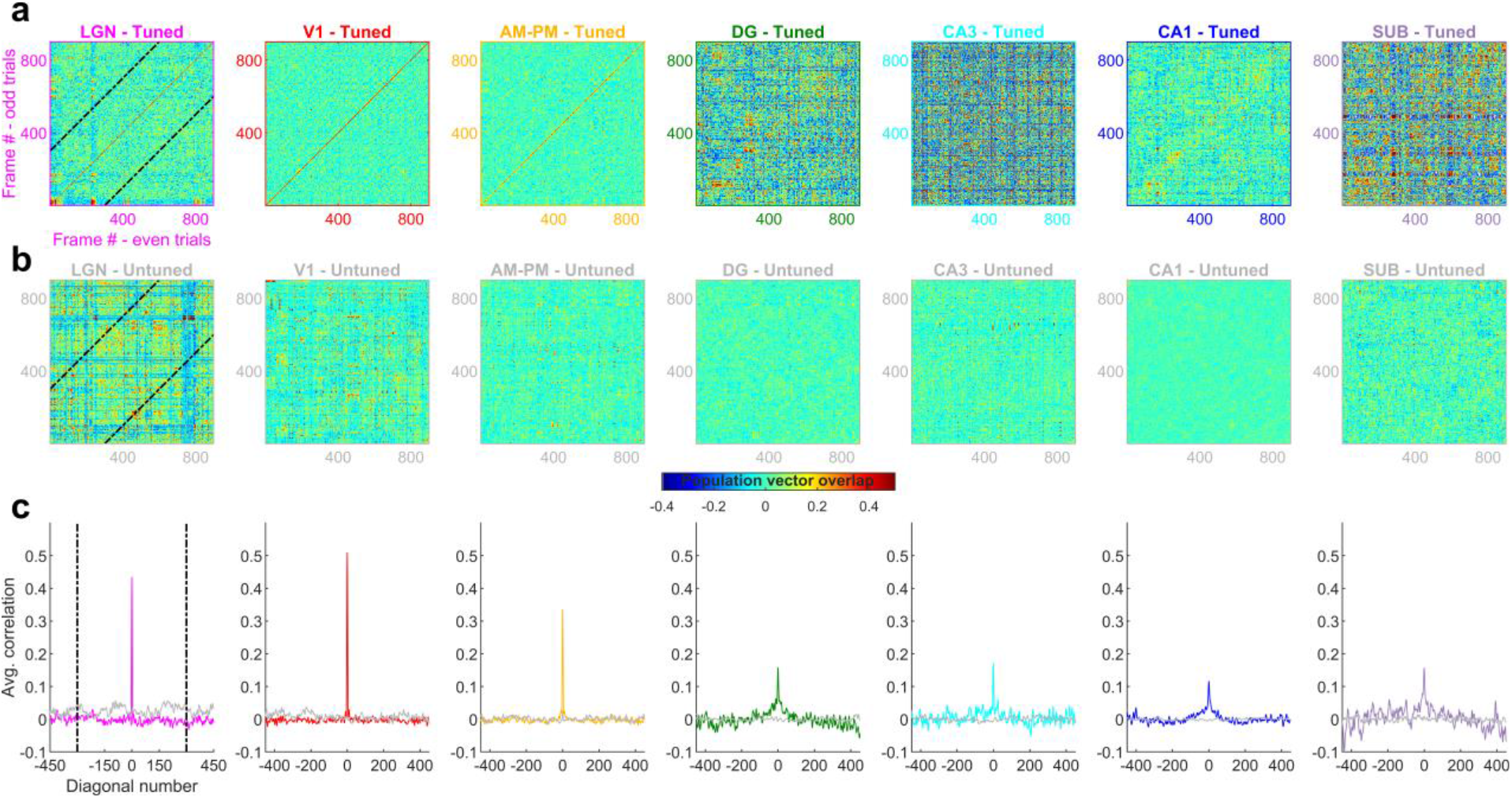
| Movie presentation did not alter hippocampal firing rates and the mega-scale coding was unrelated to cluster quality. (a) More than 50% of hippocampal place cells shut down during maze exploration^86^. In contrast, there was no consistent pattern of neural activation or shutdown during the movie presentation in all brain areas. To make a more conservative estimate, this comparison was restricted to units whose firing rates did not differ by more than 20% across the two movie blocks. Further, only the data when the animals were immobile was used to avoid confounding effects of running, and the rate threshold of 0.5Hz was removed for this panel. **(b)** The amount of movie tuning was positively correlated with the mean firing rates of the neurons for all brain regions (*r*>0.14, *p*<4.2×10^-10^). **(c)** The number of movie fields was uncorrelated with the mean firing rate of tuned cells in V1, DG, CA1 and SUB (*p*>0.12), but positively correlated for LGN, AM-PM and CA3 (*r*>0.04, *p*<0.01). Note the different y-scales for visual and hippocampal brain regions. Since the number of movie fields is an integer, data along the y-axis was slightly jittered for better visualization.**(d)** The mega-scale index was only weakly correlated with the mean firing rate of a neuron in V1 (Pearson’s correlation coefficient *r*=0.08, *p*=7.3×10^-5^), CA1 (*r*=-0.14, *p*=3.5×10^-8^) and subiculum (*r*=-0.14, *p*=0.02), and was uncorrelated for other brain regions (*p*>0.05). **(e)** The refractory violations index was uncorrelated with the mega-scale index (lower index means better cluster quality^36,108^) for all brain regions (*p*>0.05). To remove the potential confounding effect of mean firing rates, we computed the partial correlation coefficient by factoring out the mean firing rate). **(f)** Similar to (c), the isolation index (greater isolation index means better cluster quality^36,109^) was uncorrelated with the mega-scale index for all brain regions (partial correlation coefficient, by factoring out the mean firing rate, *p*>0.12). Factoring out the contribution of mean firing rate is necessary since the isolation index was typically positively correlated (the refractory violations index was typically negatively correlated) with the mean rate. The mega-scale index comparisons were restricted to movie active, tuned neurons with at least two movie peaks. Note-log spaced axes for (a)-(d)

